# CDK4 or CDK6 upregulation induces DNA replication stress and genomic instability to cause EGFR targeted therapy resistance in lung cancer

**DOI:** 10.1101/2024.03.12.584638

**Authors:** Beatrice Gini, Philippe Gui, Wei Wu, D. Lucas Kerr, Lisa Tan, Dora Barbosa, Victor Olivas, Paul Allegakoen, Carlos Gomez, Patrick R. Halliday, Sarah Elmes, Veronica Steri, Turja Chakrabarti, Trever G. Bivona, Collin M. Blakely

## Abstract

Epidermal growth factor receptor (EGFR)–mutant lung adenocarcinomas (LUAD) harbor a complex landscape of genetic co-alterations and potential oncogenic interactions. Among them are recurrent amplifications of the cell cycle regulatory genes *CDK4* and *CDK6*, which have been clinically implicated in resistance to EGFR tyrosine kinase inhibitors (TKIs). However, the mechanisms by which CDK4/6 upregulation promotes therapy resistance remain poorly defined. Here, we demonstrate that CDK4 or CDK6 overexpression limits EGFR inhibitor-induced proliferative arrest, promoting continued cell cycle progression. This is accompanied by elevated replication stress, increased TPX2 expression, DNA damage leading to ATM activation, and ultimately genomic instability. Integrative transcriptomic and copy-number analyses of EGFR-mutant LUAD tumors from both patients and preclinical models revealed that *CDK4* or *CDK6* amplification is associated with the upregulation of genes linked to tumor progression, including *AGR2*, *ASNS*, and *STEAP1*. *CDK4* amplification was also highly correlated with gene expression changes associated with epithelial-to-mesenchymal transition (EMT) in a single-cell RNA sequencing dataset from patient biopsies. In preclinical models, co-treatment with CDK4/6 and EGFR inhibitors restored proliferative arrest, induced tumor cell apoptosis, and reduced replication stress, DNA damage, and genomic instability. Our findings uncover a mechanistic basis for EGFR inhibitor resistance of *CDK4* and *CDK6* amplified EGFR-mutant LUAD. They also provide a rationale for the biomarker-driven clinical development of combination EGFR and CDK4/6-targeted therapies for the treatment of a subset of EGFR-mutant LUAD patients.

## Main Text

Targetable oncogenic driver mutations are identifiable in the majority of LUADs (1). Oncogenic activating mutations (EGFR-mutant) are present in ∼15-30% of patients in the U.S. and in more than 50% of cases in Asia (2–4). Small-molecule EGFR tyrosine kinase inhibitors (TKIs), including erlotinib, gefitinib, afatinib, dacomitinib, and third-generation EGFR inhibitors osimertinib and lazertinib, improve overall response rate (ORR) and progression-free survival (PFS) compared with traditional cytotoxic chemotherapy. However, TKI resistance limits the effectiveness of these therapies after approximately one to two years in patients with metastatic disease (5–8). Several mechanisms of acquired resistance to EGFR TKIs have been identified, including on-target mutations in *EGFR*, amplification of other receptor tyrosine kinases (RTKs), such as *MET*, oncogenic fusions involving *ALK* and *RET*, and activating mutations in the MAPK signaling pathway (9). Recent data indicate a role for genes that regulate cell cycle progression, including *CDK4* and *CDK6,* in EGFR TKI resistance (10–12).

Analysis of circulating tumor DNA (ctDNA) from a large cohort of advanced-stage EGFR-mutant LUAD patients identified co-alterations in cell cycle regulatory genes in a significant proportion of cases, which was associated with decreased response rate to osimertinib, as well as reduced PFS and overall survival (OS) (10). Among the cell cycle regulatory genes, *CDK4* and *CDK6* copy number alterations (CNAs; amplifications) were most closely associated with decreased PFS and decreased responsiveness to osimertinib treatment (10). The role of CDK4 and CDK6 in limiting osimertinib responsiveness has also emerged in preclinical studies (13–15). Recent clinical data further support the role of *CDK4* and *CDK6* gene alterations in *de novo* EGFR TKI resistance in EGFR-mutant LUAD patients (12,16). However, the mechanism(s) underlying these effects are incompletely defined.

CDK4 and CDK6 proteins regulate G1/S cell cycle progression by phosphorylating retinoblastoma protein 1 (Rb). Phosphorylation of Rb releases its inhibitory binding to transcription factor family E2F, which mediates the transition from G1 to S-phase of the cell cycle (17). While CDK4 and CDK6 have been established to limit response to oncogenic EGFR inhibition in lung cancer preclinical models and in patients, the functional characterization of the impact of CDK4/6 upregulation on TKI resistance and the mechanism(s) underlying these effects have not been clearly defined beyond an association with changes in Rb phosphorylation and the G1/S cell cycle transition. Alterations in cell cycle genes have been shown to induce deregulation of cell cycle checkpoints, compromising DNA integrity and contributing to the accumulation of DNA damage and abnormal chromosomal segregation (18–20). Here, we provide evidence that CDK4 or CDK6 upregulation induces DNA replication stress and genomic instability in EGFR-mutant LUAD, activating TPX2 and ATM, which are known to promote EGFR TKI resistance (21,22). In addition, we identified recurrent copy-number changes associated with CDK4 and CDK6 upregulation and increased expression of genes related to EMT, which may also contribute to tumor progression. Co-treatment with a CDK4/6 inhibitor reversed many of these effects and re-sensitized EGFR-mutant LUAD to EGFR TKI treatment in both *in vitro* and *in vivo* patient-derived models. These findings provide novel mechanistic and therapeutic insights into EGFR-mutant LUAD.

## Results

### CDK4 or CDK6 upregulation promotes sustained DNA synthesis and cell-cycle progression during EGFR inhibition in EGFR-mutant LUAD

To investigate the mechanism(s) by which CDK4 or CDK6 may promote decreased responsiveness of EGFR-mutant lung cancer cells to EGFR TKI treatment, we utilized patient-derived EGFR-mutant LUAD cell lines: H1975 (EGFR^L858R/T790M^, CDK4/6^wt^) and PC9 (EGFR^del19^, CDK4/6^wt^); a primary, patient-derived organoid (PDO), TH107 (EGFR^del19^, CDK4/6^wt^); and a patient-derived xenograft (PDX), TH116 (EGFR^L858R^, CDK4^wt^, CDK6^AMP^) (Supplementary Tables 1-2). We overexpressed CDK4 or CDK6 in H1975 and PC9 EGFR-mutant LUAD cells (Supplementary Fig. 1A-D) and quantified protein expression. This demonstrated a similar degree of CDK4 overexpression observed in HCC827 (EGFR^del19^, CDK4^AMP^, CDK6^wt^) cells with endogenous *CDK4* amplification and CDK6 overexpression to a similar degree observed in the TH116 PDX (EGFR^L858R^, CDK4^wt^, CDK6^AMP^) (Supplementary Fig. 1E-J), indicating a pathophysiology-relevant degree of protein overexpression. CDK4 and CDK6-overexpressing PC9 and H1975 cells demonstrated relative resistance to osimertinib in colony formation assays (Supplementary Fig. 2). TH107 (EGFR^del19^, CDK4/6^wt^) PDOs, which had no evidence of *CDK4* or *CDK6* CNAs (Supplementary Fig. 3A, B), were genetically engineered to overexpress CDK4 or CDK6 through lentiviral infection to model *CDK4/6* CNAs (Supplementary Fig. 3C) (23). This resulted in decreased responsiveness to osimertinib treatment in multiple assays (Supplementary Fig. 3D-H). These results confirm that CDK4 or CDK6 overexpression confers relative resistance to the EGFR TKI osimertinib across multiple preclinical models of EGFR-mutant LUAD.

To assess the effects of CDK4 or CDK6 overexpression on EGFR signaling in the context of TKI treatment *in vivo*, we assessed how an osimertinib dose range modulates EGFR phosphorylation and signaling in CDK4 or CDK6-overexpressing H1975 CDXs (Fig. 1A). Osimertinib (at 5 mg/kg/day) was sufficient to inhibit EGFR signaling in E.V. control and CDK4 or CDK6-overexpressing CDXs (Fig. 1A, B). However, CDK4 or CDK6 overexpression conferred relative resistance to osimertinib treatment compared to E.V. control H1975 CDXs (Fig. 1C, D, and Supplementary Fig. 4) and in CDK6 overexpressing PC9 CDXs (Supplementary Fig. 5).

**Figure 1.**
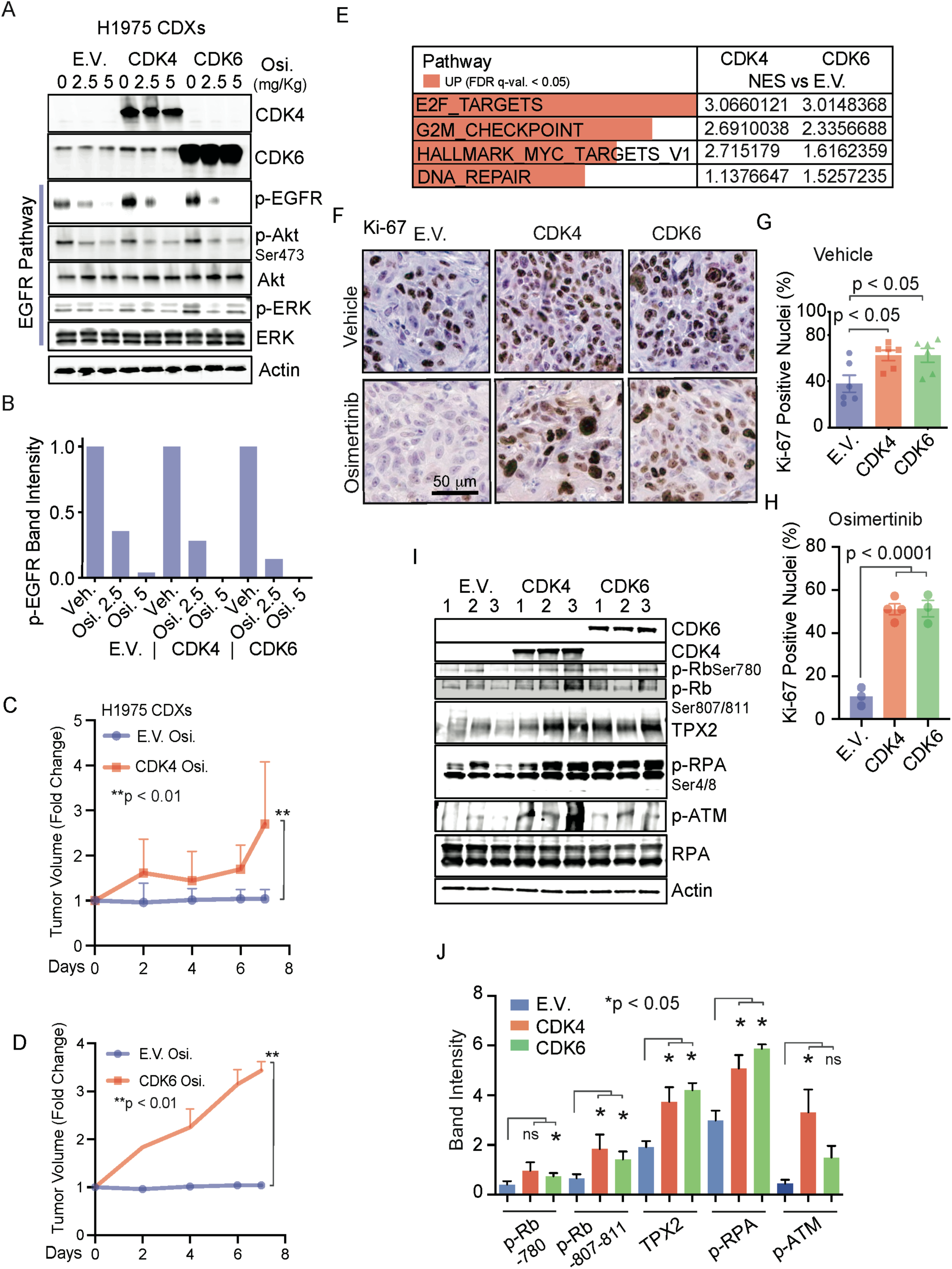
CDK4/6 upregulation promotes osimertinib resistance and replication stress in EGFRmt LUAD CDXs. (A) Immunoblot analysis of CDK4/6 overexpressing and empty vector (E.V.) H1975 EGFR-mutant LUAD CDXs treated with vehicle (0 mg/kg) or osimertinib (2.5-5 mg/kg). Mice were treated by oral gavage for 4 days and sacrificed 4 hours after the final dose. Protein expression detected by immunoblotting as indicated. (B) Quantification of p-EGFR immunoblotting is shown in (A). (C, D) Growth curves of osimertinib (5 mg/Kg) treated CDK4 (C) or CDK6 (D), overexpressing H1975 CDXs, compared to empty vector controls (n = 4-6 xenografts per group; error bars representing SEM). (E) Gene expression differences were derived from RNA-sequencing analysis of vehicle-treated xenografts. Top significantly up-(red) regulated GSEA pathways (n = 3 xenografts per group; p-value calculated as described in methods; NES: Normalized Enrichment Score). (F-H) Representative IHC-stained images (F) and quantification (G, H) of Ki-67 from E.V. and CDK4/6 overexpressing H1975 CDXs treated with osimertinib (5 mg/Kg) (n = 3-6 xenografts per group; p-value assessed by one-way ANOVA and Tukey’s multiple comparisons test; error bars representing SEM). (I, J) Immunoblotting of replication stress biomarkers p-Rb, TPX2, p-RPA, and p-ATM in CDK4/6 overexpressing H1975 CDXs (I) and quantification (J) versus empty vector (E.V.) controls (error bars representing SEM; following a significant ANOVA result (p < 0.05), post-hoc analysis was conducted using Tukey’s Honest Significant Difference (HSD) test for pairwise comparisons. Statistical significance is indicated by brackets and p-values, with a threshold of p < 0.05. (E.V.: empty vector; CDK4 and CDK6: CDK4 or CDK6 overexpressing CDXs; Veh.: vehicle; Osi.: osimertinib).

To assess which downstream pathways may be affected by CDK4 and CDK6 activation, we evaluated gene expression changes in CDXs overexpressing CDK4 and CDK6 compared with controls. Gene set enrichment analysis (GSEA) of CDK4 or CDK6 overexpressing H1975 CDXs showed significant upregulation of E2F1 target genes, which mediate G1/S transition (17), together with substantial enrichment of G2/M checkpoint, HALLMARK_MYC_TARGETS_V1, and DNA repair genes (Fig. 1E and Supplementary Fig. 6A-D). These data suggest that EGFR-mutant LUAD CDXs overexpressing CDK4 or CDK6 upregulate the E2F1 transcriptional program, which promotes G1/S transition, entry in the S-phase of the cell cycle, and proliferation (17,24).

We further evaluated this by quantifying the expression of the proliferation biomarker Ki-67, which was significantly higher in the CDK4- and CDK6-overexpressing tumor xenografts than in controls and was maintained upon treatment with osimertinib (Fig. 1F-H). The quantity of newly synthesized DNA, measured by EdU incorporation in CDK4- and CDK6-overexpressing tumor xenografts, was also increased compared to controls in both the vehicle and osimertinib treatment groups (Supplementary Fig. 7A-C). These data demonstrate that overexpression of CDK4 or CDK6 results in increased DNA synthesis and cellular proliferation maintained during osimertinib treatment.

### CDK4 or CDK6 upregulation leads to DNA replication stress in EGFR-mutant LUAD

We hypothesized that increased DNA synthesis and proliferation observed in *in vivo* models could lead to DNA replication stress and, potentially, replication stress-associated DNA damage in CDK4- and CDK6-overexpressing LUADs. We evaluated biomarkers indicative of DNA replication stress and DNA damage repair (DDR) (25,26), including the phosphorylation of Rb protein, expression of TPX2, a microtubule nucleation factor, and phosphorylation of replication protein A (RPA). CDK4 and CDK6-expressing CDXs demonstrated higher expression levels of these DNA replication stress biomarkers than controls (Fig. 1I, J, and Supplementary Fig. 7D, E).

Given the role of TPX2 during interphase as a protective factor of DNA fork stability during replication stress (26) and its role in EGFR TKI resistance (22), we assessed the nuclear expression of TPX2 in CDK4 and CDK6-overexpressing H1975 CDXs by immunohistochemistry. We identified increased TPX2 nuclear expression compared to controls (Supplementary Fig. 7F, G). Increased TPX2 nuclear localization in CDK4 and CDK6 overexpressing CDXs was also observed post-osimertinib treatment, although it was decreased compared with pre-treatment levels (Supplementary Fig. 7F, G). *TPX2* gene expression has been shown to depend on the activity of transcription factor E2F1 (26). E2F1 knockdown resulted in the downregulation of *TPX2* in EGFR-mutant NSCLC cells (Supplementary Fig. 7H, I). Together, these results demonstrate that CDK4 and CDK6 overexpression induce biomarkers of DNA replication stress in EGFR-mutant lung cancer *in vivo* models.

DNA replication stress can also induce DNA damage and activation of DNA damage response (DDR) pathways, as the G1/S checkpoint is necessary to ensure cellular repair of DNA damage before DNA replication (27). We assessed phosphorylation of ataxia-telangiectasia mutated (ATM) as a DDR biomarker and known promoter of EGFR TKI resistance (21), as well as γ-H2AX and EdU double immunofluorescence staining in CDK4 and CDK6-overexpressing tumor xenografts as markers of DNA damage in proliferating cells. CDK4 and CDK6-overexpressing CDXs demonstrated increased expression of phosphorylated ATM (Fig. 1I, J), indicative of DDR activation. Further analysis of RNA sequencing data from CDK4- and CDK6-overexpressing H1975 CDXs revealed significant upregulation of key DDR inducers and effectors, including *CHEK1*, *CHEK2*, and *RAD51* (Fig. 2A, B), supporting this hypothesis. CDK4 and CDK6-overexpressing H1975 CDXs also showed significantly higher γ−H2AX foci in proliferating cells than controls (Fig. 2C, D). This effect persisted during osimertinib treatment in CDK4- and CDK6-overexpressing tumors (Fig. 2E, F). These data highlight the accumulation of DNA replication stress and damage upon CDK4 and CDK6 upregulation, which persist upon osimertinib treatment in EGFR-mutant LUAD cells.

**Figure 2.**
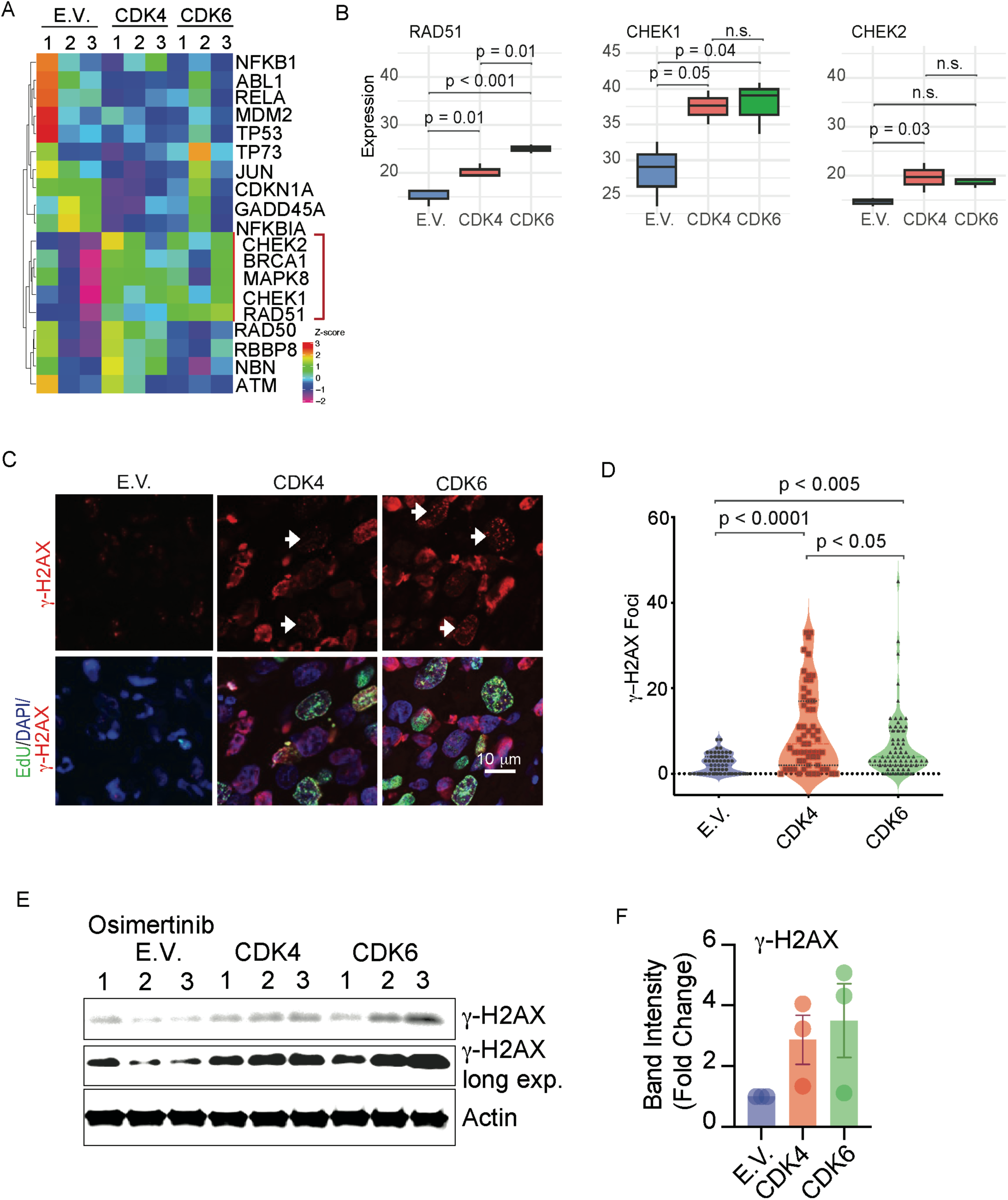
CDK4/6 upregulation promotes DNA damage accumulation in EGFRmt LUAD CDXs treated with osimertinib. (A, B) Gene expression differences were derived from RNA-sequencing analysis of vehicle-treated xenografts. For the vehicle- and osimertinib-treated groups, mice received daily oral gavage for 7 days and were sacrificed 4 hours after the final dose. Gene expression profiling of empty vector (E.V.), CDK4, and CDK6 overexpression H1975 CDXs demonstrating statistically significant enrichments in the ATM pathway genes *RAD51*, *CHEK1*, and *CHEK2*, across CDK4/6 overexpressing CDXs compared to E.V. controls (n = 3 xenografts per group; p-value assessed by one-way ANOVA and Tukey’s multiple comparisons test). (C, D) Immunofluorescence stain for γ-H2AX (red) and EdU (green), with DAPI stain (blue) (C) and quantification of γ-H2AX foci number (D) in EdU-positive cells for vehicle-treated H1975 CDXs (white arrows point to double-positive cells; n= 50-100 EdU-positive cells per sample, one xenograft per group; p-value calculated with one-way ANOVA and Tukey’s multiple comparisons test). (E, F) Immunoblot analysis of γ-H2AX expression in CDK4/6 overexpressing H1975 CDXs treated with osimertinib for one week compared to E.V. overexpressing control CDXs (E) and blot’s quantification (F) (n = 3 xenografts per group; fold change compared to E.V. controls; error bars representing SEM). (E.V.: empty vector; CDK4 and CDK6: CDK4 or CDK6 overexpressing CDXs; Veh.: vehicle; Osi.: osimertinib).

### CDK4 or CDK6 upregulation increases genomic instability and the acquisition of tumor-promoting CNAs in EGFR-mutant LUAD

Given the replication stress and DNA damage observed with CDK4 and CDK6 upregulation in preclinical models, we investigated whether amplification or overexpression of CDK4 or CDK6 drives genomic instability in EGFR-mutant LUAD. As a measure of genomic instability, we assessed the fraction of genome altered (FGA) using whole-exome sequencing of CDK4- and CDK6-overexpressing EGFR-mutant NSCLC CDXs treated with either vehicle or osimertinib. We observed a 15- to 20-fold increase in FGA in CDK4 and CDK6-overexpressing tumors compared to E.V. controls under both conditions (Fig. 3A), suggesting that CDK4 or CDK6 up-regulation increases genomic instability. Despite the elevated structural GIN, tumor mutation burden (TMB) remained unchanged across CDK4, CDK6, and E.V. xenografts in both vehicle and osimertinib treatment arms, as determined by SNV analysis (Supplementary Fig. 8A, B), suggesting that CDK4/6-associated genomic instability primarily involves copy number alterations rather than increased point mutations. Integration of RNA-seq with CNA data revealed a subset of genes that showed both CN gains and transcriptional upregulation in CDXs overexpressing CDK4 (Fig. 3B) and CDK6 (Fig. 3C). Several known pro-tumor effectors, including *AGR2*, *CBX3*, *ANLN*, *SEC61G*, *PSMA2*, *STEAP1*, and *GGCT,* were significantly upregulated at both the DNA and RNA levels in both CDK4 and CDK6 overexpression CDXs (Fig. 3B-E) (27–34). Notably, these overlapping CNA and RNA expression signatures persisted upon osimertinib treatment (Fig. 3F), suggesting that upregulation of these genes may contribute to EGFR TKI resistance. Gene ontology and pathway enrichment analysis of the overlapping genes revealed strong enrichment for pathways involved in cell cycle regulation, DNA replication, and DNA repair, consistent with the role of CDK4 and CDK6 in driving proliferation and managing replication stress-induced DNA damage (Supplementary Fig. 8C, D). These results indicate that upregulation of CDK4 and CDK6 is associated with increased FGA and transcriptional activation of DNA repair and cell-cycle regulatory genes, which may contribute to genomic instability and resistance to EGFR inhibition.

**Figure 3.**
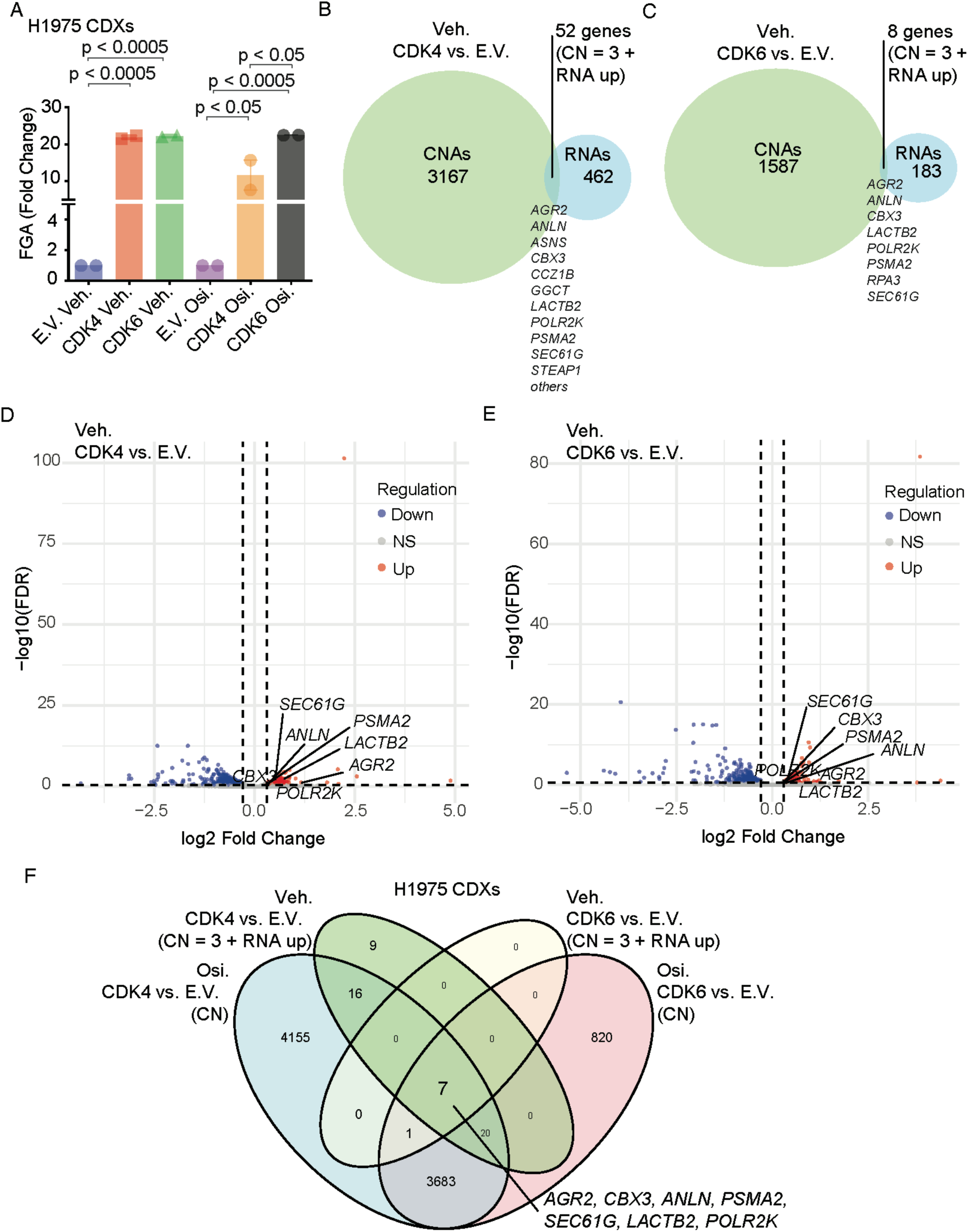
Overlap of CDK4/6 upregulated genes with abnormal CN alterations in CDK4/6 overexpressing EGFRmt LUAD CDXs. (A) Fraction of Genome Alterations (FGA) analysis from whole exome sequencing data from osimertinib (5 mg/kg) or vehicle-treated H1975 CDK4/6 overexpressing CDXs compared to empty vector (E.V.) controls. Mice were treated by daily oral gavage for 7 days and sacrificed 4 hours after the final dose (n = 2 xenografts per group; p-value assessed by one-way ANOVA and Tukey’s multiple comparisons test; error bars representing SEM). (B, C) Venn diagram illustrating the overlap between genes upregulated in CDK4 (B) or CDK6 (C)-overexpressing CDXs (RNA-seq) and genes with copy number alterations (CNAs) in the vehicle-treated arm (normalized to E.V. CDXs). RNA-seq upregulated genes are shown in sky blue, and CNA-associated genes are shown in green. The intersection indicates genes both transcriptionally upregulated and genomically altered. (D, E) Volcano plot illustrating differential gene expression between CDK4 (D) or CDK6 (E)-overexpressing CDXs vs. E.V. controls. Genes were classified as upregulated (red), downregulated (blue), or not significant (grey) based on log2 fold change (FC) > 0.3 and FDR < 0.3. Selected genes of interest (*AGR2, CBX3, ANLN*, etc.) are annotated. Vertical and horizontal dashed lines represent fold-change and FDR thresholds, respectively. (F) Venn diagram showing the overlap of genes between osimertinib-treated samples (Osi.) and vehicle-treated samples (Veh.), based on upregulated RNA expression (RNA up) and/or copy number alterations (CN = 3; normalized to E.V. CDXs). (CNAs detected in CDK4 and CDK6 overexpressing CDXs normalized to empty vector control E.V. CDXs in the same treatment group; CDK4: CDK4 overexpressing CDXs; CDK6: CDK6 overexpressing CDXs; for the CNAs analysis, n = 2 xenografts for each group; for the RNAseq experiments, n = 3 xenografts for each group).

### CDK4 or CDK6 inhibition reduces DNA replication stress and genomic instability in CDK4/6 upregulated EGFR-mutant CDXs and PDXs

Combination therapy with osimertinib and the CDK4/6 inhibitor palbociclib has been shown to inhibit Rb phosphorylation, induce cell-cycle arrest, and decrease tumor cell survival in EGFR-mutant cell lines (15). Similarly, we found that combination treatment with osimertinib and palbociclib was sufficient to induce regression in CDXs overexpressing CDK4 or CDK6 (Supplementary Fig. 9A-D). To assess whether a reduction in DNA replication stress and genomic instability accompanied these effects, we measured Ki-67, p-Rb, p-RPA, and γ-H2AX expression in tumors treated with osimertinib alone, palbociclib alone, or in combination. Osimertinib plus palbociclib-treated tumors exhibited decreased Ki-67 expression (Fig. 4A, B), as well as reduced expression of p-Rb, p-RPA, and γ-H2AX compared to tumors treated with either drug alone (Fig. 4C-F). Osimertinib plus palbociclib-treated tumors also exhibited increased apoptosis, as evidenced by increased expression of cleaved caspase-3 (Supplementary Fig. 9E, F). The percentage of CDK4-positive cells was also decreased in osimertinib plus palbociclib-treated tumors (Fig. 4G, H), suggesting that CDK4/6 inhibition may eliminate cells with CDK4 upregulation and concomitant CDK4-induced DNA replication stress and genomic instability. Overall, these findings demonstrate that combined EGFR and CDK4/6 inhibition reduces proliferation, DNA replication stress, and genomic instability associated with CDK4 or CDK6 upregulation, thereby increasing tumor cell apoptosis and tumor regression.

**Figure 4.**
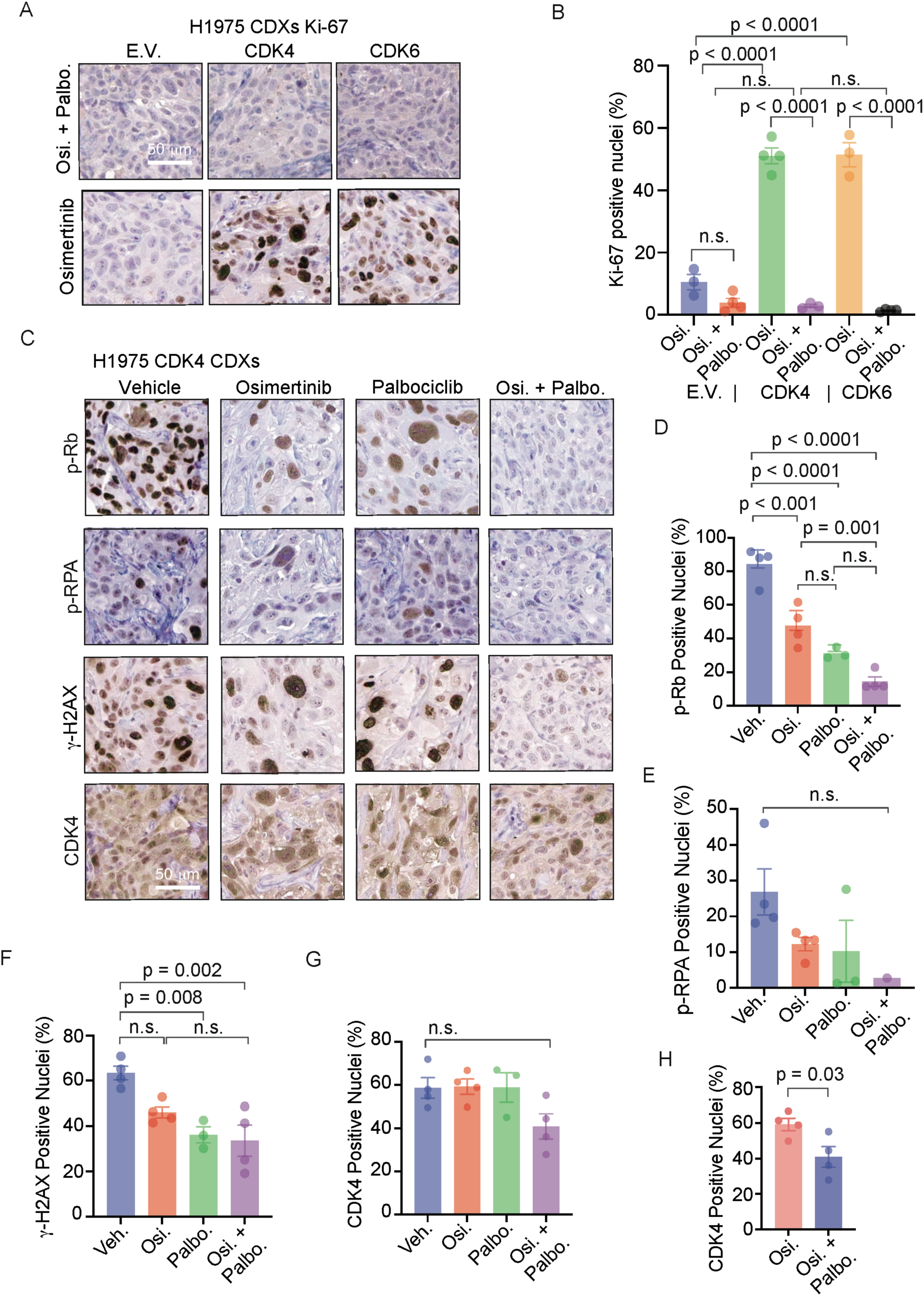
Combination therapy with EGFR and CDK4/6 inhibitors suppresses tumor growth and decreases replication stress and genomic instability in CDK4/6 overexpressing EGFRmt LUAD CDXs (A, B) Representative Ki-67 IHC-stained tumor sections. (A) and quantification (B) from E.V. and CDK4/6 overexpressing H1975 CDXs treated with osimertinib (5 mg/kg) or combinatorial treatment with palbociclib (100 mg/kg). Mice were treated by daily oral gavage for 7 days and sacrificed 4 hours after the final dose (n = 3-4 xenografts per group; p-value assessed by one-way ANOVA and Tukey’s multiple comparisons test; error bars representing SEM). (C-H) Representative IHC stained tumor sections (C) and quantification of p-Rb (D), p-RPA (E), γ-H2AX (F), and CDK4 (G, H) from CDK4 overexpressing H1975 CDXs treated with osimertinib (5 mg/kg) and combinatorial treatments with palbociclib (100 mg/kg) (n = 3-4 xenografts per group for Ki-67, p-Rb, p-RPA, γ-H2AX, p-RPA IHC stains; n = 1 for p-RPA in the combinatorial treatment group due to high necrosis in the tissues; p-value assessed by one-way ANOVA and Tukey’s multiple comparisons tests and Student’s T-test; error bars representing SEM). (E.V.: empty vector; CDK4: CDK4 overexpressing CDXs; CDK6: CDK6 overexpressing CDXs; Veh.: vehicle; Palbo.: palbociclib; Osi.: osimertinib; p-value assessed by one-way ANOVA and Tukey’s multiple comparisons test; error bars representing SEM).

We extended these findings to primary EGFR-mutant PDX (Supplementary Fig. 1) and PDO (Supplementary Fig. 3) models harboring concurrent *CDK6* amplification or overexpression. We included abemaciclib, another CDK4/6 inhibitor, to verify that the observed effects are consistent across different CDK4/6 inhibitors. In line with the CDK4-overexpressing CDX data, we found that combination treatment with osimertinib and abemaciclib was sufficient to induce regression of CDK6 overexpressing primary EGFR-mutant PDOs (Supplementary Fig. 10A). To investigate whether *CDK6* amplification induces DNA replication stress and genomic instability that can be overcome by CDK4/6 inhibition in a more clinically relevant model, we utilized a PDX model (TH116) developed from a LUAD patient biopsy at resistance to osimertinib treatment, which harbored concurrent *EGFR* p.L858R mutation and *CDK6* amplification (Supplementary Fig. 1E, F, and Supplementary Tables 1-2). EGFR-mutant, *CDK6*amp TH116 PDXs demonstrated continued growth during osimertinib treatment, indicating EGFR TKI resistance (Supplementary Fig. 10B, C). Similar to CDK6 overexpressing CDX models, these *CDK6*amp, EGFR-mutant PDXs maintained high p-Rb phosphorylation when treated with osimertinib at doses sufficient to suppress EGFR and ERK phosphorylation (Fig. 5A). The PDXs also demonstrated high levels of the DNA replication stress and DNA damage markers TPX2, p-AURKA, in the osimertinib-treated tumors (Fig. 5A). γ-H2AX was also elevated in vehicle and osimertinib-treated tumors, consistent with DNA damage (Fig. 5B, C). An enrichment of CDK6-positive nuclei was observed upon osimertinib treatment, suggesting that EGFR inhibition may select for cells with upregulated nuclear CDK6 (Fig. 5B, D, E). Treatment of TH116 PDXs with palbociclib, either alone or in combination with osimertinib, significantly reduced both γH2AX levels as well as FGA (Fig. 5C, F) and decreased tumor growth more than osimertinib alone (Supplementary Fig. 10C-F). The combinatorial treatment of osimertinib with palbociclib resulted in the highest degree of FGA reduction, although not statistically significant compared to monotherapy with osimertinib (Fig. 5F). Mutant Allele Frequency (MAF) and TMB were also assessed to evaluate overall genomic burden, but neither showed significant changes across treatment groups (Supplementary Fig. 10G, H).

**Figure 5.**
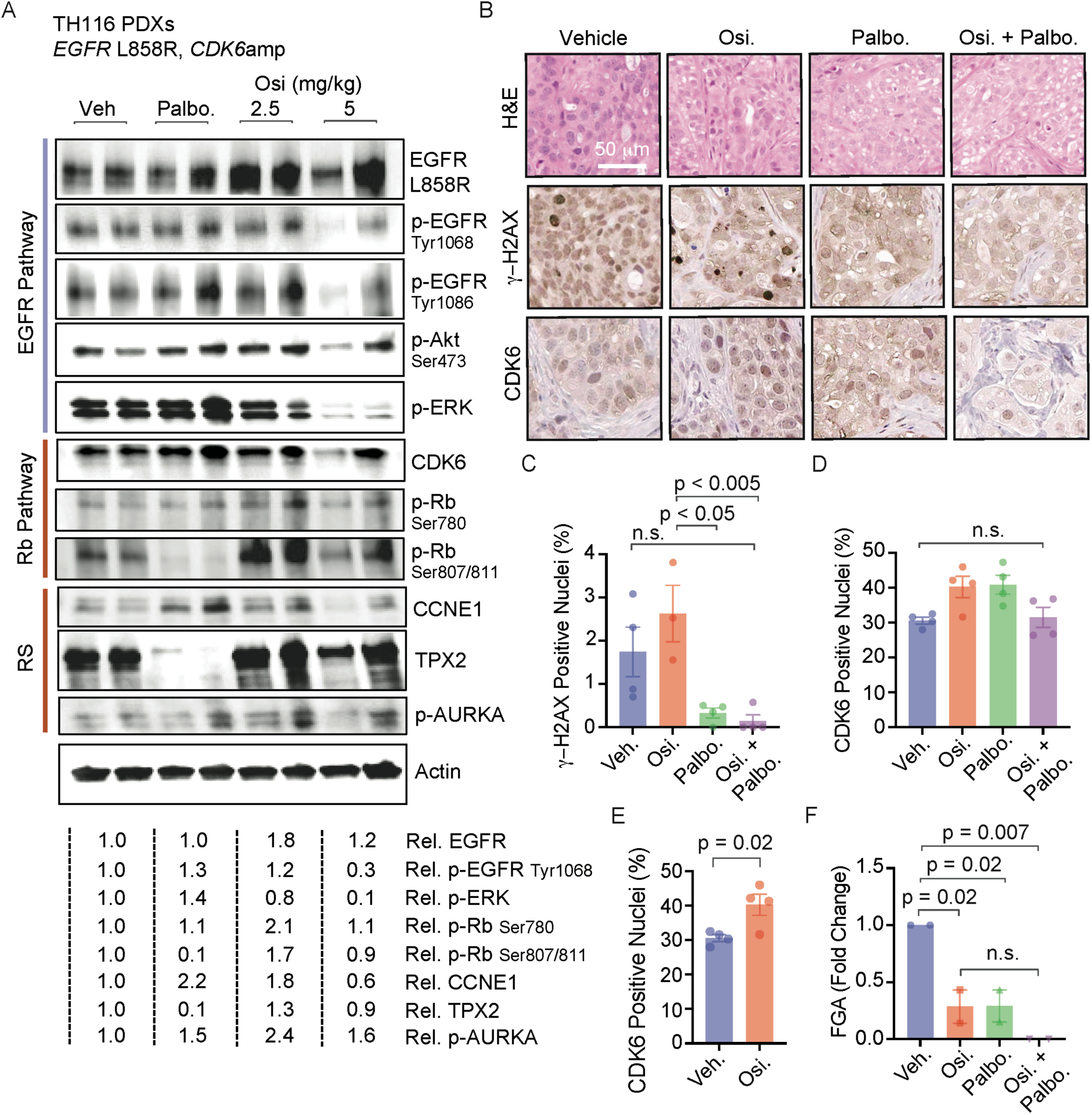
Combination therapy with EGFR and CDK4/6 inhibitors decreases replication stress and genomic instability in a *CDK6*amp, EGFRmt primary LUAD PDXs. (A) Immunoblotting with quantification for EGFR pathway, CDK6-Rb pathway, and replication stress biomarkers CCNE1, TPX2, and p-AURKA from TH116 (EGFR L858R, *CDK6amp)* primary patient-derived xenografts (PDXs), treated with vehicle, palbociclib (150 mg/kg), or osimertinib (5 mg/kg) by oral gavage for 4 days and sacrificed 4 hours after the final dose. Normalized quantification of the level of a panel of biomarkers is provided in the lower panel. (B-E) Representative hematoxylin/eosin, γ-H2AX, and CDK6 IHC-stained tumor sections (B) and quantification (C-E) using tissues from TH116 PDXs (n = 4 xenografts per group; Student’s T-test). (F) FGA analysis in TH116 PDXs, treated with vehicle, osimertinib (5 mg/kg), palbociclib (100 mg/kg), or a combination of osimertinib and palbociclib (Osi. + Palbo.). Mice were treated by oral gavage for 38 days and sacrificed 4 hours after the final dose (n = 2 xenografts per group; error bars representing SEM.Veh.: vehicle; Palbo.: palbociclib; Osi.: osimertinib; p-value assessed by one-way ANOVA and Tukey’s multiple comparisons tests; error bars representing SEM).

Overall, these results suggest that combinatorial treatment with osimertinib and a CDK4/6 inhibitor in CDK4/6-upregulated and CDK6-amplified EGFR-mutant LUAD CDX and PDX models is effective and significantly reduces DNA replication stress and genomic instability, as evidenced by decreased expression of DNA replication stress and DNA damage markers (γ-H2AX, p-RPA), reduced Ki-67 and p-Rb levels, and lower FGA.

### *CDK4* or *CDK6* copy-number alterations are associated with significantly increased genomic instability in EGFR-mutant LUAD patient tumors

To extend our preclinical findings and assess the clinical relevance of CDK4 and CDK6 CNAs in EGFR-mutant LUAD, we analyzed two large clinical datasets: Foundation Medicine (n = 660) and MSK-IMPACT GENIE (n = 1983), comprising a combined total of 2,643 EGFR-mutant advanced LUAD cases (Supplementary Table 3). Utilizing targeted exome sequencing data across 401 cancer-relevant genes, Foundation Medicine cases were stratified into two groups: tumors with cell cycle gene alterations, based on TCGA definitions, and those without (Supplementary Fig. 11A, B). Among cell cycle-altered tumors, frequent events included deletions or mutations in tumor suppressors *CDKN2A* (49.0%), *CDKN2B* (38.7%), and *RB1* (21.8%), followed by CNAs in *CDK4* (15.7%), *CCNE1* (10.1%), *CCND1* (8.1%), and *CDK6* (6.7%). *CDK4* and *CDK6* alterations were significantly enriched in EGFR-mutant versus KRAS-mutant LUAD (Supplementary Fig. 11C), were mutually exclusive from one another (Supplementary Fig. 11D, E), and predominantly represented by copy number gains (Supplementary Fig. 11F, G) in the MSK-IMPACT GENIE and Foundation Medicine clinical cohorts. To evaluate the relationship between *CDK4/6* CNAs and genomic instability, we quantified the FGA and TMB as surrogate markers. EGFR-mutant LUAD harboring cell cycle gene alterations exhibited significantly elevated FGA and TMB relative to cell cycle-negative tumors across both datasets (Fig. 6A, B, and Supplementary Fig. 12A, B). Subgroup analysis further revealed that tumors with *CDK4* or *CDK6* CNAs had markedly increased FGA (Fig. 6C, D). Tumors with *CDK4* CNAs showed increased FGA in the MSK-IMPACT, but not the Foundation Medicine dataset (Fig. 6E, F), whereas those with *CDK6* CNAs demonstrated increased FGA in both datasets (Fig. 6G, H). FGA correlated with the degree of copy number gain of *CDK4* or *CDK6* (Fig. 6I). Notably, no corresponding increase in TMB was observed with *CDK4* or *CDK6* CNA, similar to our findings in preclinical models (Supplementary Fig. 12C–H). *CDK6* CNAs were associated with higher FGA compared to *CDK4* CNAs in the Foundation Medicine dataset (Supplementary Fig. 12I), but not the MSK-IMPACT (Supplementary Fig. 12J). *RB1*-altered tumors also demonstrated an increase in FGA compared to *RB1*-non-altered tumors (Fig. 6J); however, the vast majority of tumors with *CDK4/6* CNAs were not found to have *RB1* alterations (Fig. 6K, L). CNAs in *CCND1*, *CCND3*, *CDKN2A*, *CDKN2B*, or *CCNE1* exhibited minimal impact on FGA (Supplementary Fig. 13A–E). Importantly, *TP53* mutations were evenly distributed between cell cycle-positive and cell cycle-negative cohorts (68.3% vs. 62.7%, p = 0.13; Supplementary Fig. 11A, B), suggesting that *TP53* status alone did not account for the increased FGA. To determine whether increased FGA was specific to *CDK4/6*-related alterations, we evaluated tumors with *MDM2* and *NFKBIA* amplifications, commonly observed in EGFR-mutant LUAD. Neither *MDM2* nor *NFKBIA* amplification was associated with increased FGA (Supplementary Fig. 13F, G), highlighting the specificity of *CDK4/6* amplification on FGA. Next, we assessed whole genome doubling (WGD)—a hallmark of genomic instability—using TCGA WES data. Tumors harboring *CDK4* or *CDK6* amplification demonstrated significant enrichment for WGD events (Supplementary Fig. 13H–K), supporting the conclusion that *CDK4* or *CDK6* amplification contributes to genomic instability.

**Figure 6.**
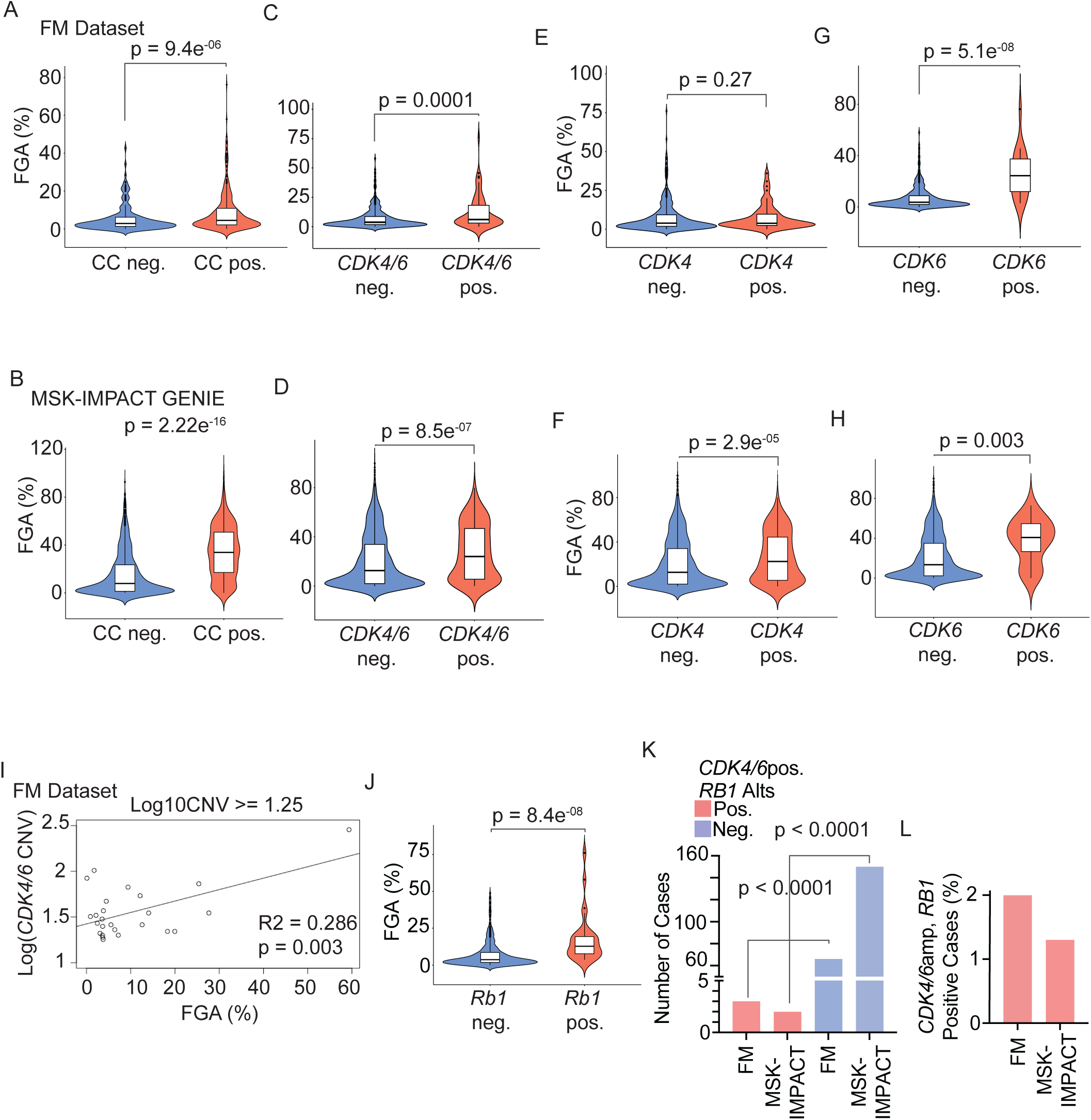
Concurrent *EGFRmt* and *CDK4/6* CNAs are associated with an enhanced percentage of genome alterations in LUAD patient tumors. Fraction of Genome Alterations (FGA) analysis in EGFRmt LUAD using Foundation Medicine (FM) (A) or MSK-Impact GENIE (B) targeted exome sequencing datasets. (A) FGA was compared in tumors harboring concurrent cell cycle CNAs (CC pos.) to tumors negative for such alterations (CC neg.) in the FM (A) and MSK-Impact GENIE datasets (B). (C, D) FGA analysis in EGFRmt tumors stratified by *CDK4/6* CNA status in the FM dataset (C) and MSK-Impact GENIE dataset (D). (E, F) FGA analysis in EGFRmt tumors stratified by *CDK4* CNA status in the FM (E) or MSK-Impact GENIE (F) datasets. (G, H) FGA analysis in EGFRmt tumors stratified by *CDK6* CNA status in the FM (G) or MSK-Impact GENIE (H) datasets. (I) Positive correlation plot showing degrees of *CDK4/6* amplification (y-axis) and corresponding FGA (p-value calculated with F-statistics) (FM Dataset; each dot representing one *CDK4/6* pos. tumor). (J) FGA analysis with FM dataset comparing *RB1* alteration positive versus *RB1* alteration negative EGFRmt LUAD cases. (K, L) Mutual exclusivity analysis (K) and percentage of cases (L) carrying concurrent *CDK4/6* amp and *RB1* gene alterations (SNVs and CNAs) in EGFRmt tumors from Foundation Medicine (FM) or MSK-Impact GENIE targeted exome sequencing (p-value calculated with Fisher’s test). (FGA boxplots included in violin plots represent the interquartile range, lower quartile 25%, median, and upper quartile 75%; p-value calculated with Wilcoxon test).

### Integrative single-cell and bulk RNA sequencing analyses reveal *CDK4/6* amplification**‒**associated transcriptional reprogramming

To determine whether the gene expression changes identified in CDK4/6-overexpressing H1975 CDXs are also present in patient tumors, we analyzed bulk RNA-seq data from the OncoSG EGFR-mutant LUAD cohort. *CDK4/6*-amplified (n = 9) were compared to *CDK4/6* non-amplified tumors (n = 84). Patient tumors showed significantly higher expression of six genes̶*AGR2*, *ASNS*, *CCZ1B*, *GGCT*, *PSMA2*, and *STEAP1* that were previously found to be upregulated in the CDX model (Fig. 3 and Supplementary Fig. 14). To verify these findings in an independent cohort, we analyzed single-cell RNA sequencing data from a cohort of EGFR-mutant LUAD clinical samples collected at our institution (35). This cohort included three tumors with *CDK4* amplification and 11 tumors from patients without *CDK4* or *CDK6* amplification, as determined by clinical sequencing and confirmed by whole-exome sequencing. No tumors with *CDK6* amplification were identified. UMAP visualizations showed higher cancer cell–specific CDK4 expression in *CDK4*-amplified tumors compared with *CDK4/6*wt tumors (Fig. 7A-D). Significantly higher expression of *AGR2*, *ASNS*, *GGCT*, *STEAP1*, and *CCZ1B,* but not *PSMA2,* was also observed in *CDK4*-amplified compared to non-amplified tumors (Fig. 7E and Supplementary Fig. 15A, B). Spearman correlation analyses revealed positive correlations between *CDK4* expression and *STEAP1*, *AGR2*, and *ASNS* in cancer cells from *CDK4*-amplified tumors compared to non-amplified tumors (Fig. 7F). These genes have been implicated in promoting tumor progression across multiple cancer types (29–31). In addition to higher expression of these cancer-promoting genes, *CDK4*-Amplified tumors exhibited significantly higher epithelial-mesenchymal transition (EMT) scores than non-amplified tumors (Fig. 7G). These findings suggest that *CDK4* amplification may promote transcriptional programs, including EMT, associated with aggressive tumor behavior.

**Figure 7.**
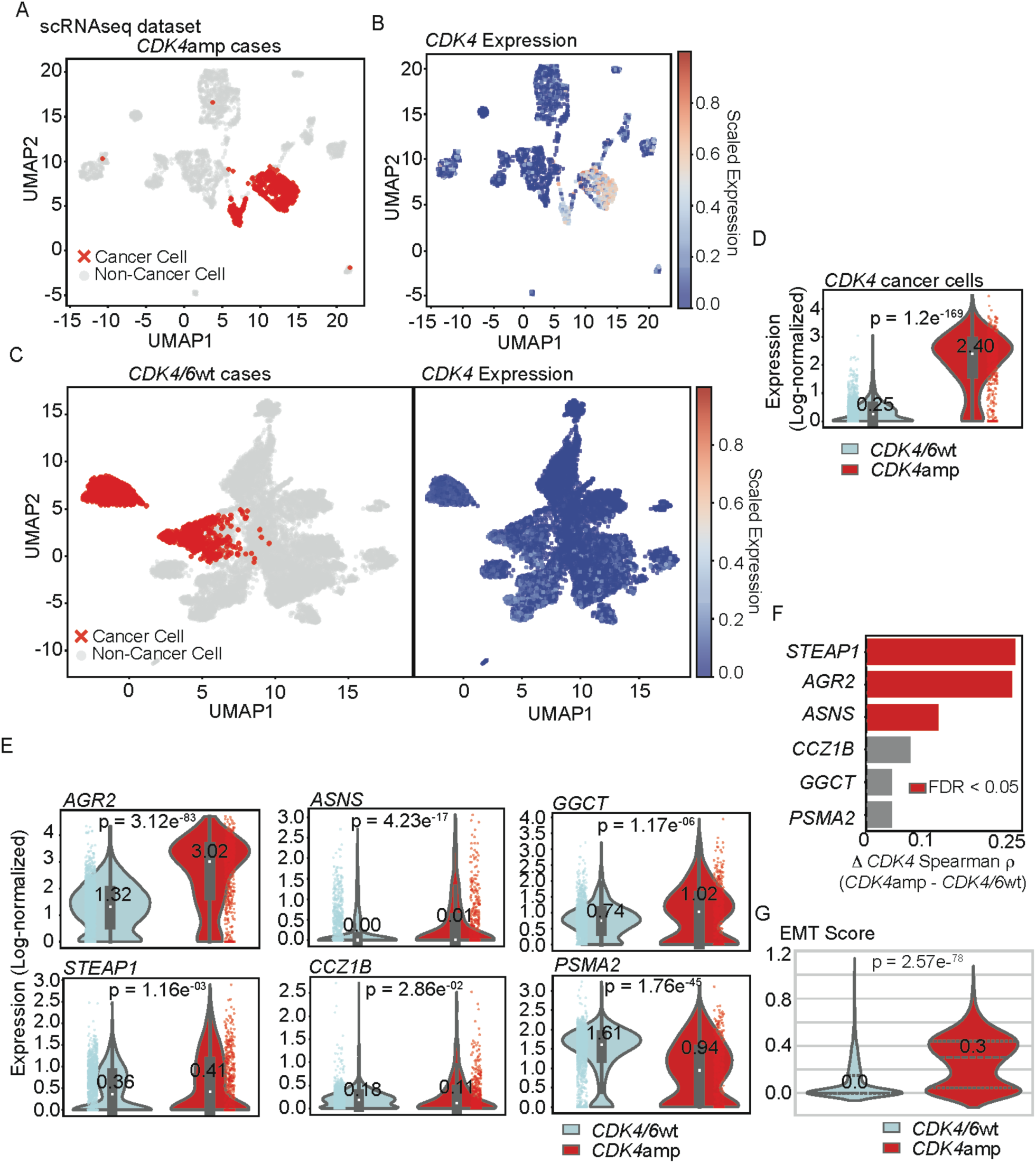
Transcriptional changes associated with *CDK4*amp in EGFRmt LUAD cancer cells from patient biopsies. (A) UMAP visualization of single-cell RNA-seq data from *CDK4*amp, EGFRmt LUAD clinical samples (n = 3), with cancer cells annotated in red based on inferCNV inference. (B) UMAP showing high *CDK4* expression in *CDK4*amp tumor cells, demonstrating localization of CDK4 elevated expression to cancer cells. (C) UMAP visualization of single-cell RNA-seq data from *CDK4/6*wt EGFRmt LUAD clinical samples (n = 11, left scatterplot) and UMAP showing a low degree of *CDK4* expression across *CDK4/6*wt tumor cells (right scatterplot). (D) Violin plots comparing *CDK4* expression in cancer cells from *CDK4*amp (n =3) versus *CDK4/6*wt (n = 11) tumors, showing significantly higher *CDK4* expression in *CDK4*amp cases. (E) Violin plots pointing to *AGR2*, *STEAP1*, *GGCT*, and *ASNS* expression in cancer cells as significantly upregulated in *CDK4*amp versus *CDK4/6*wt tumors. (F) Difference in Spearman correlation of gene expression with *CDK4* gene expression between single cancer cells from *CDK4*amp (n = 483) and *CDK4/6*wt (n = 3188) EGFR-mutant LUAD patients (Δρ). P-values were calculated by comparing Fisher z-transformed correlations using a z-test, then adjusted for multiple testing using the False Discovery Rate (FDR). Bars in red indicate genes with significant differences after FDR correction (FDR < 0.05); gray bars are not significant. (G) Violin plot comparing epithelial-mesenchymal transition (EMT) scores in single cancer cells from *CDK4*amp (n = 483) and *CDK4/6*wt (n = 3188) EGFR-mutant LUAD patients. EMT scores were calculated as the mean expression of canonical EMT markers (*VIM*, *ZEB1*, *SNAI1*, *TWIST1*, *CDH2*, and *FN1*). Medians are indicated above each violin. Statistical comparison was performed using a two-sided Mann–Whitney U test.

## Discussion

This study provides a mechanistic framework for how complex somatic genetic interactions shape the biology and therapeutic vulnerabilities of oncogene-driven lung adenocarcinoma (Fig. 8). By combining genomic analyses with preclinical models and patient-derived xenografts, we demonstrate that amplification of *CDK4* or *CDK6*—occurring in approximately 10% of EGFR-mutated lung cancer cases—can drive DNA replication stress, genomic instability, and resistance to EGFR tyrosine kinase inhibitors.

**Figure 8.**
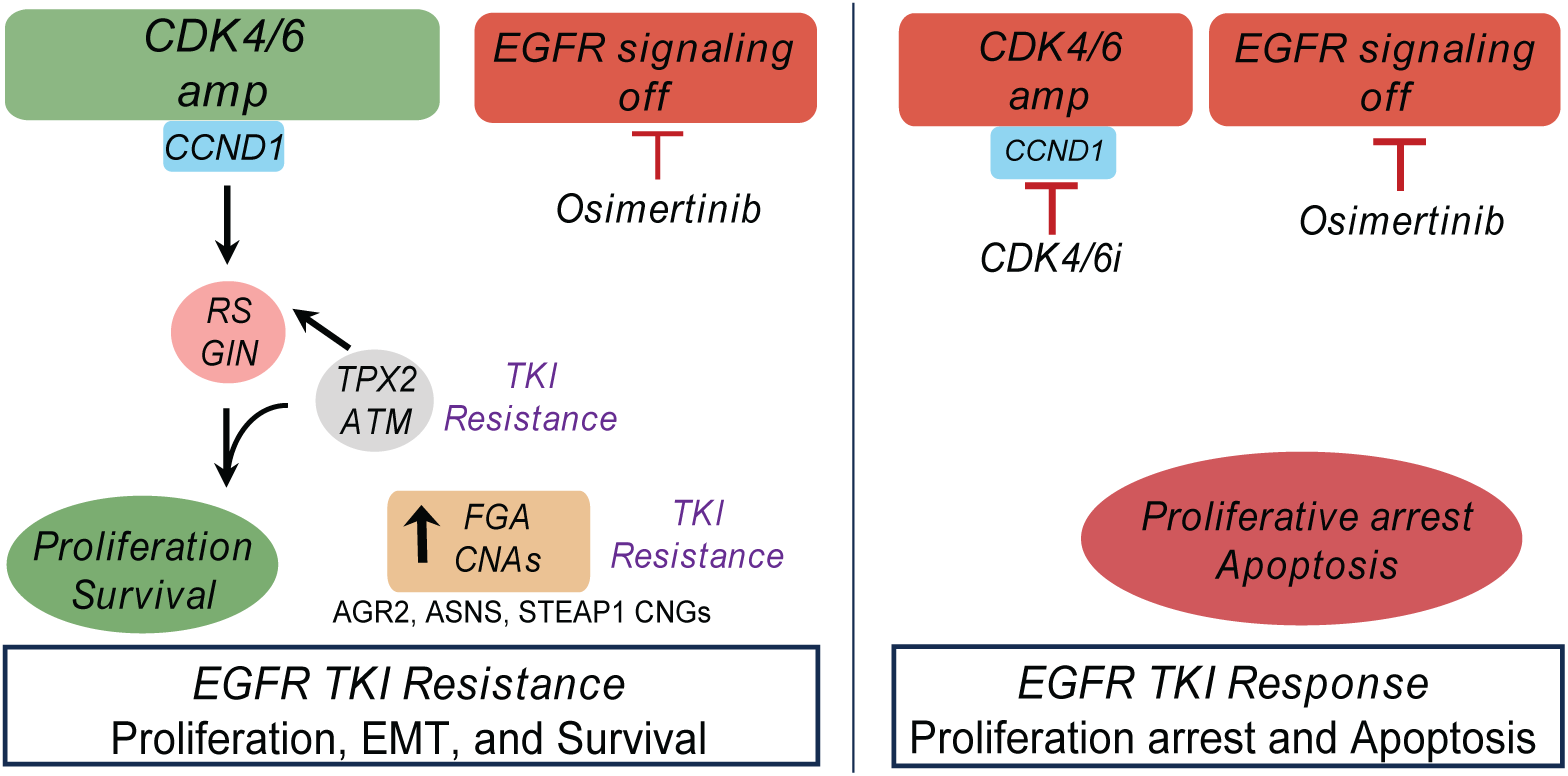
Model of the effects of *CDK4* or *CDK6* amplification in EGFRmt LUAD. (Left) Schematic model illustrating the impact of *CDK4* or *CDK6* amplification in EGFRmt LUAD. In *CDK4*amp or *CDK6*amp tumors, Rb phosphorylation persists despite EGFR TKI treatment, leading to continued proliferation, accumulation of replication stress (RS) and genomic instability (GIN), and activation of TKI resistance mechanisms (e.g., TPX2/AURKA, ATM). This promotes an increased fraction of genome altered (FGA) and genomic and transcriptional upregulation of tumor promoting genes (e.g., *AGR2*, *ASNS*, *STEAP1*), and epithelial to mesenchymal transition (EMT) contributing to EGFR TKI resistance. (Right) Co-treatment with a CDK4/6 inhibitor (e.g. palbociclib or abemaciclib restores G1/S checkpoint control, reduces proliferation, and induces apoptosis, leading to tumor regression.

We show that *CDK4/6* amplification is associated with an increased fraction of genome altered (FGA) in two large, independent patient cohorts (Foundation Medicine and MSK-IMPACT GENIE), linking copy-number gains to chromosomal instability. These findings are clinically significant, as increased FGA has been associated with poor prognosis and recurrence in NSCLC (36). Similarly, in the TRACERx study, increased copy-number heterogeneity was associated with genomic instability and an increased risk of recurrence or death (37,38). Interestingly, other cell cycle alterations, such as *CCNE1* amplification or *CDKN2A/2B* deletions, were not associated with increased FGA, underscoring the specificity of the CDK4/6 effect. Moreover, *CCNE1* amplification was mutually exclusive with *CDK4/6* amplification, suggesting distinct and non-redundant roles in tumor biology. Notably, alterations across a broader number of cell cycle genes were associated with increased TMB in addition to increased FGA. This likely reflects the fact that broad cell cycle alterations included genes whose loss of function is known to affect DNA repair, including *CDKN2A/B* and *RB1*. In contrast, CDK4/6 amplification primarily promoted replication stress and chromosomal instability, but is not known to impact DNA repair, resulting in increased copy-number alterations (FGA) without a corresponding increase in TMB.

Mechanistically, *CDK4/6* amplification overrides the G1 arrest typically induced by EGFR TKI treatment (39), thereby enabling continued entry into S-phase and proliferation. In both treatment-naïve and osimertinib-treated xenografts, *CDK4/6* overexpression correlated with sustained p-Rb signaling, increased EdU and Ki-67 labeling, and upregulation of DNA replication stress and DNA damage response markers, including TPX2, γ-H2AX, p-RPA, and p-ATM.

Upregulation of TPX2, a transcriptional target of E2F (24,26), is associated with AURORA Kinase A activation and resistance to EGFR inhibitors (22), suggesting a potential downstream mechanism by which *CDK4/6* amplification promotes EGFR TKI resistance. Similarly, ATM activation has been associated with EGFR-mutant lung cancer cell survival and EGFR TKI resistance (21). Activation of TPX2-AURKA and ATM in response to replication stress and DNA damage may thus represent two mechanisms by which *CDK4/6* amplification promotes EGFR TKI resistance (Fig. 8).

In the context of promoting cell cycle progression, replication stress, and DNA damage, CDK4 and CDK6 activation led to recurrent copy number changes of *AGR2*, *ASNS*, and *STEAP1* in both preclinical models, patient-derived xenografts, and in patient tumors. Intriguingly, *ASNS* and *AGR2* have been associated with WNT/β-catenin activation (29,40), while *AGR2* and *STEAP1* have been associated with EMT (31,40). WNT/β-catenin and EMT have each been associated with EGFR TKI resistance (10,41,42), suggesting additional mechanisms by which *CDK4/6* amplification contribute to poor response to EGFR TKI treatment. This suggests that while CDK4/6 hyperactivation induces broad structural genomic instability, it also leads to recurrent amplification and transcriptional upregulation of specific pro-tumor loci including *AGR2*, *ASNS*, and *STEAP1*. This enrichment likely reflects a multi-step evolutionary process rather than purely localized genomic fragility or direct E2F-driven upregulation. In this model, CDK4/6-mediated replication stress establishes a background of random structural variations, but subsequent positive selection under EGFR inhibitor pressure drives the clonal expansion of cells harboring copy number gains at these specific loci. Because AGR2, ASNS, and STEAP1 are associated with aggressive phenotypes like EMT and WNT/β-catenin activation, the enrichment of cancer cells harboring these alterations may allow them to overcome the suppressive effects of EGFR inhibitor treatment. Indeed, we found that *CDK4-*amplified EGFR-mutant lung cancer cells exhibited increased expression of genes associated with EMT compared with *CDK4*-non-amplified cancer cells in single-cell RNA-sequencing analyses of patient tumors.

This finding resonates with recent evidence that *RB1* loss promotes reversible lineage plasticity and accelerates EGFR TKI resistance by enabling resistant mesenchymal states (43). Together, these observations point to a convergent role of the CDK4/6-RB axis in promoting transcriptional plasticity and EMT, highlighting that therapeutic resistance in lung cancer may be shaped as much by adaptive cell-state transitions as by proliferative signaling.

To overcome the pro-tumor effects of *CDK4/6* amplification, we explored combinatorial therapy targeting EGFR and CDK4/6. Combined osimertinib and CDK4/6 inhibitor treatment suppressed proliferation, reduced DNA damage and chromosomal alterations, and induced apoptosis in EGFR-mutant tumors with *CDK4* or *CDK6* amplification or overexpression. This highlights a potential therapeutic strategy to delay or prevent resistance, which should be further evaluated in clinical trials among biomarker-selected patients. A phase II trial of osimertinib and the CDK4/6 inhibitor abemaciclib has been completed; however, results are not yet publicly available. This trial was open to patients regardless of *CDK4/6* amplification status. Based on our data, we predict that biomarker selection for patients with *CDK4/6* amplification will be necessary to observe meaningful clinical benefit.

Our study has limitations. We focused primarily on the canonical cell-cycle functions of CDK4/6, while emerging roles in immune modulation, metabolism, and transcriptional regulation remain unexplored in this context (44). Here, we also highlight a link to EMT programs in single-cell analyses, where only *CDK4*-amplified tumors were available, suggesting that *CDK4* amplification may promote resistance through transcriptional reprogramming toward mesenchymal phenotypes. Further analyses will be needed to determine whether *CDK6* amplification has a similar effect. Additionally, sequencing approaches limited the detection of low-frequency subclones and non-coding regulatory alterations that may contribute to the evolution of resistance. Future studies will be needed to address these dimensions across broader cohorts.

In summary, we uncovered multiple mechanisms of EGFR inhibitor resistance in LUAD driven by CDK4/6 hyperactivation. CDK4/6 hyperactivation promotes continued cell proliferation during EGFR TKI treatment, leading to DNA replication stress and DNA damage, which in turn activates TPX2 and ATM, known drivers of EGFR TKI resistance. DNA replication stress and damage, in turn, lead to chromosomal instability with increased copy-number changes, which include copy-number gains in genes associated with EMT-associated transcriptional plasticity (Fig. 8). These findings establish a rationale for therapeutic strategies combining EGFR and CDK4/6 inhibitors to suppress primary oncogenic signaling and prevent replication stress and DNA damage and their downstream consequences. This also has potential therapeutic implications for other cancers that harbor co-activation of oncogenes within the RTK-RAS-MAPK pathway, concurrent with cell cycle-promoting genomic alterations, which occur in a high proportion of epithelial malignancies. More broadly, our work illustrates how co-occurring genomic alterations and transcriptional programs synergistically shape tumor evolution and therapy response.

## Methods

### Cell Lines, Primary Organoids and Reagents

All cell lines were obtained and cultured according to the American Type Culture Collection (ATCC) recommendations. Before testing, cell lines were authenticated through STR profiling and confirmed negative for mycoplasma contamination. EGFR mutant (EGFRmt) LUAD cell lines used in the present study were H1975 (EGFR L858R, T790M), HCC827 (EGFR Del19), and PC9 (EGFR Del19). EGFR-mutant cells were cultured in RPMI 1640 medium (HyClone, GE Healthcare), containing 10% FBS (SAFC, Sigma-Aldrich) and 1X penicillin and streptomycin (HyClone, GE Healthcare). DMEM (HyClone, GE Healthcare) supplemented with 10% FBS, 0.1X penicillin, and streptomycin was used for HEK293-FT cells. All cell lines were cultured in a humidified incubator with 5% CO_2_ at 37 °C. Primary, patient-derived organoid TH-107 cells were established from a biopsy specimen and cultured in Matrigel as described previously(45,46). Drugs used in these studies were Osimertinib, Abemaciclib (Selleckchem), and Palbociclib (LC Laboratories). The antibodies used for immunoblotting, immunohistochemistry, and immunofluorescence were: CDK4 (Cell Signaling Tech., 12790) CDK6 (Cell Signaling Tech., 3136), pEGFR-Tyr1086 (ThermoFisher, 369700), p-EGFR-Tyr1068 (Cell Signaling Tech., 3777), EGFR L858R (Cell Signaling Tech., 3197), p-AURKA (Cell Signaling Tech., 3079), p-Akt (Cell Signaling Tech., 4060), p-ERK (Cell Signaling Tech., 4370), Actin (Sigma, A2228), Rb (Cell Signaling Tech., 9309), p-Rb-Ser780 (Cell Signaling Tech., 9307), p-Rb-Ser807/811 (Cell Signaling Tech., 8516), CCNE1 (Cell Signaling Tech., 20808), TPX2 (Cell Signaling Tech., 12245), cleaved caspase 3 (Cell Signaling Tech., 9661), p-RPA (Bethyl Laboratories, A300 245A M), p-ATM (Cell Signaling Tech., 13050), RPA (Bethyl Laboratories, A300 244A), γ-H2AX (EMD Millipore, 05-636), ki67 (Leica, NCL-L-Ki67-MM1). All antibodies were diluted per the datasheet recommendations. Small interfering RNAs for E2F1 and scramble control were purchased from Dharmacon, and transient knockdown was induced following the manufacturer’s instructions.

### Lentiviral Infection, Soft Agar, and Crystal Violet Assays

Mammalian expression vectors EGFP, CDK4, and CDK6 were purchased from Origene. Transfection of HEK293-FT cells with the overexpressing vectors was performed using Mirus reagent, per the manufacturer’s instructions. Viral particles were collected after 72 hrs of transfection to infect EGFR-mutant LUAD cells in a medium containing polybrene (8 μg/ml; Sigma-Aldrich). After 72 hrs, cells were sorted for EGFP or selected with puromycin (1 μg/mL; Gibco). Selected cells were used in the in vitro and in vivo tests. Successful overexpression of CDK4 and CDK6 was confirmed by qPCR and/or immunoblotting.

In vitro, soft-agar colony formation assay was performed as previously described(47). Colonies’ growth was assessed for up to 4 weeks in the presence of osimertinib (10-100 nM) or DMSO control. At the time of collection, 6-well soft agar plates were incubated with a solution of 0.005% crystal violet in water for three hours. Plates were then imaged using ImageQuant (GE Healthcare), and colonies were counted using the ImageJ program (NIH).

### Preclinical Studies

H1975, PC9 xenografts, and TH-116 PDXs were established in compliance with the UCSF IACUC IRB-approved protocols. Each H1975 and PC9 flank xenograft was obtained after subcutaneous injection of 1 million cells in each 4-week-old female SCID or athymic mouse flank. NSG mice were used for the propagation and establishment of TH-116 PDXs. Osimertinib was reconstituted for in vivo experiments following previously published work (48). Osimertinib treatments were performed by daily oral gavage (dose range assessed: 2.5 mg/Kg, 5 mg/Kg, 10 mg/Kg). Mice euthanasia was done when xenografts reached 20 mm in diameter. We used similar drug regimens for combination studies. Mice were randomly divided and kept unlabeled until the treatments started. Multiple researchers measured the tumors with a digitized caliper.

IP injections of 100 μL EdU (200 μg) solution in PBS were carried out 48 hrs before xenografts resection (Click-iT EdU kit, Invitrogen)(49) to assess the degree of newly synthesized DNA. OCT blocks of tissues were collected from each tumor and used for deriving cryosections that were stained either following the manufacturer’s protocol for single EdU stain (Click-iT EdU kit, Invitrogen) or double immunofluorescence stain with γH2AX (50).

### Immunoblotting, Immunohistochemistry, Immunofluorescence

Protein lysates were extracted in RIPA buffer, adding protease inhibitors (Roche) and phosphatase inhibitors (Roche). For the immunoblotting, 15 μg of proteins were loaded into precast 4–15% acrylamide gels (Bio-Rad), then transferred to nitrocellulose membranes with the Trans-blot Turbo Transfer system (Bio-Rad). Blots were blocked for 1 hr at room temperature in Tris-buffered saline, 0.1% Tween20 (vol/vol), and 5% (vol/vol) BSA (Fisher Scientific). Incubation with the primary antibodies was performed overnight at 4 °C, after which the membranes were washed twice with Tris-buffered saline, Tween20 (0.1% vol/vol). Incubation with secondary HRP-conjugated antibodies (Cell Signaling Technology, anti-rabbit IgG, no. 7074, anti-mouse IgG, no. 7076) was run for 1 hr at room temperature. Membranes were then incubated with ECL reagent (GE Healthcare), and the signal was detected by chemiluminescence. The development and scanning of the blots were run with ImageQuant LAS4000 (GE Healthcare Life Sciences). ImageJ (NIH) allowed western blot quantification.

Formalin-Fixed, Paraffin-Embedded tissue blocks and sections were provided by the UCSF histology core. Immunohistochemistry (IHC) was performed as previously described (10). The stained slides were scanned with Aperio ScanScope CS Slide Scanner (Aperio Technologies) using a 20x objective. ScanScope algorithm was applied for the IHC quantification, considering 3-5 fields per section and calculating mean, and S.E.M. CDK4, CDK6, p-Rb, γ-H2AX, p-RPA stains were quantified on the same areas of adjacent xenograft, viable tissue sections.

FFPE sections were stained with hematoxylin-eosin for the tissue necrotic area analysis, and necrosis was quantified as the percentage of affected areas following pathological guidelines (51).

### Bioinformatics Analysis of CDK4/6-Overexpressing H1975 Xenografts

#### Fraction of Genome Alteration Analysis (FGA), SNVs, and copy number calling from whole exome sequencing data from in vivo tumor tissues

DNA was extracted from xenograft tissues using a tissue-DNA extraction-specific kit (Qiagen) and sent to Novogene (Sacramento, USA) for whole exome sequencing (WES; NovaSeq 6000, PE150, 200X). Pair-end .fastq files were mapped to the hg19 genome. Mutation (SNVs) calling was done using the SeqMule pipeline. The .vcf files were annotated using ANNOVAR software at a high-performance computing cluster (UCSF Helen Diller Comprehensive Cancer Center). Further analysis of annotated variants was conducted under the RStudio/R environment. Cell-derived and patient-derived xenograft WES bam files were used to infer copy number alterations between the treatment and control groups, utilizing the “copy number” function in the VarScan algorithm. The altered genomic segments were annotated at the gene level. Each gene length was retrieved from hsapiens.UCSC.hg19.knownGene database. The sum of altered CNV genes over the size of captured WES was defined as a Fraction of Genome Alterations (FGA). We normalized the CDK4/6 overexpressing and CDK6 amp xenografts FGA with the FGA from empty vector, CDK4/6 wild type, or vehicle-treated control xenografts expressing it as a fold-change (52). To infer the absolute integer gene copy number, mapped BAM files were used as input along with hg19_SureSelect_Human_All_Exon_V5.bed and human_g1k_v37.fasta files for CNVkit (default parameters, python library, https://cnvkit.readthedocs.io/en/stable/), and the log2 ratio of estimated segmentation between two conditions was computed to identify regions significantly affected by CNVs.

### RNA sequencing and GSEA

Total RNAs were extracted from CDK4/6 overexpressing tissues and empty vector controls (Qiagen) and sent to Novogene for RNA sequencing (NovaSeq 6000, PE150).

For the Gene Set Enrichment Analysis (GSEA), a gene expression matrix for CDK4 and CDK6 overexpressing tissues, and controls (cells expressing empty vector, E.V.) in triplicate was imported into the GSEA algorithm (v4.03, Broad Institute). MSigDB was used for gene set enrichment comparisons between CDK4/6 overexpressing and E.V. CDXs groups. The enrichment score (ES), normalized ES, and False discovery rate (FDR) were provided by the algorithm. An FDR < 25% was used for exploratory enrichment analysis.

### Integration of Gene Expression and Copy Number

Differential Gene Expression Analysis (Vehicle Group) RNA-seq differential expression results were obtained from CDK4-, CDK6-, and EGFP (E.V.)-overexpressing H1975-derived xenografts in the vehicle-treated group. Gene-level log₂ fold changes and FDR-adjusted p-values were provided for CDK4 vs. EGFP and CDK6 vs. E.V. comparisons. Differentially expressed (DE) genes were identified using relaxed criteria: |log₂ fold change| ≥ 0.3 and false discovery rate (FDR) ≤ 0.3. Volcano plots were generated to visualize differential gene expression. Genes were color-coded based on significance and direction of change: upregulated (red), downregulated (blue), or not significant (grey). Selected genes of interest (e.g., AGR2, CBX3, ANLN) were annotated for interpretability. DE analyses were conducted before and after Xengsort filtering, which removed approximately 6% mouse reads to assess potential bias from murine stromal contamination. Agreement between filtered and unfiltered data was quantified using: • Pearson and Spearman correlation of log₂ fold change values • Fisher’s exact test for overlap significance • Jaccard index for DEG set similarity. These comparisons were implemented in Python v3.10 using matplotlib, seaborn, and matplotlib-venn. Results demonstrated near-perfect consistency (e.g., CDK4 DE: Pearson r = 1.000, Jaccard = 0.991), confirming minimal impact of mouse filtering on DE calls. Copy number variants (CNVs) were inferred from whole-exome sequencing (WES) data using CNVkit, with normalization performed relative to the EGFP (E.V.) control group. CNVkit-derived results were summarized as log₂-transformed copy number ratios and estimated absolute copy number values per gene. Log₂ ratios reflect relative copy number changes, while absolute CN values were inferred and rounded by CNVkit for discrete copy number calling. Gene CNA lists were derived from the CDK4 and CDK6 upregulated CDXs by annotating the genes in the chromosomal segments affected by CNAs (UCSC Genome Browser).

Venn diagrams were constructed to identify genes concurrently upregulated at both transcript and copy number levels to visualize overlap between DE genes (RNA-seq) and CN-amplified genes (CNVkit) in the vehicle-treated group.

Pathway Enrichment Analysis Genes overlapping between transcriptomic (vehicle group) and copy number (vehicle group) datasets were converted to Entrez IDs using the bitr() function from the clusterProfiler package (with org.Hs.eg.db as the reference). Functional enrichment was performed via: • Gene Ontology (GO): enrichGO • KEGG pathways: enrichKEGG • Reactome pathways: enrichPathway from ReactomePA. All enrichment results were adjusted using the Benjamini-Hochberg FDR correction, with significance defined as adjusted p < 0.05. Results were visualized using barplots, dotplots, and enrichment maps.

### Foundation Medicine, MSK-IMPACT GENIE Clinical Dataset Analyses

EGFR-mutant advanced-stage LUAD cases analyzed from Foundation Medicine (FM) were 660, and targeted exome sequencing data for a panel of 401 cancer-related genes were used to detect somatic mutations and copy number alterations. Foundation Medicine provided the annotated data matrix. FGA analysis was computed from the Foundation Medicine dataset using a panel of 306 cancer-related genes with CNA (gains and deletions), following previously published methods (52). Briefly, patients were stratified based on the presence of selected cell cycle gene copy number alterations (53). For each patient, genes with copy number alterations annotated from Foundation Medicine were filtered in, and the size of each gene was retrieved. The sum of altered gene sizes over the captured genome was computed as FGA. Boxplots were created with interquartile range (lower quartile 25%, median, and upper quartile 75%). Each dot represents the FGA value in each patient’s tumor. Statistical significance was assessed using the Wilcoxon test. For the Tumor Mutation Burden (TMB) analysis, the total nonsynonymous mutations over the captured genome size were computed as tumor mutation burden (TMB, number of gene mutations/Mega-base). Cases with concurrent *EGFR* and *CDK4/6* alterations were further analyzed for *RB1* and *TP53* alteration frequencies.

FGA analysis was computed from the MSK-IMPACT GENIE dataset using 1983 EGFR-mutant LUAD cases stratified based on the presence of selected cell cycle gene copy number alterations (53) as previously described (AACR project GENIE Consortium, GENIE Cohort version 15.1 – public). Cases with concurrent *EGFR* and *CDK4/6* alterations were further analyzed for *RB1* and *TP53* alteration frequencies.

For the correlation analysis between FGA and *CDK4/6* amplifications, patients harboring *CDK4/CDK6* amplifications with copy numbers greater than 4 were selected, and the value of log10(CNV CDK4/6 amp) versus FGA value was plotted with a linear regression model for the best-fit line, and the F-statistic was used to calculate the p-value.

Oncoprint analysis was used to show the most frequent concurrent gene alterations in cell cycle (CC) alteration-positive (n=357) or cell cycle-alteration-negative (n=303) Foundation Medicine dataset cohorts. Each column represents a patient; each row represents one gene. Gene alteration frequencies greater than 6% were shown in the oncoprints.

Analysis of LUAD cases through cBioPortal was done by selecting cases positive for CDK4/6 CNAs with concurrent EGFR mutations (125 total cases).

### Whole Genome Doubling (WGD) analysis in the TCGA LUAD dataset

Whole Genome Doubling (WGD) analysis from the TCGA lung adenocarcinoma (LUAD) cohort dataset was downloaded from: https://github.com/judithabk6/ITH_TCGA/blob/master/external_data/TCGA_mastercalls.abs_tables_JSedit.fixed.txt

Correlation analysis was run to annotate WGD events along with the status of *CDK4/6* amp using the TCGA CNA file annotations. Fisher’s exact test was used to measure the statistical significance.

### Gene Expression and Statistical Analysis in OncoSG EGFR mutant Cohort

Z-score normalized gene expression data were analyzed for EGFR-mutant LUAD cases in the OncoSG cohort. CDK4/6-positive (n = 9) and CDK4/6-negative (n = 84) groups were compared using Welch’s two-sample t-test. Significance thresholds were set at *P* < 0.05. Violin plots were generated in *ggplot2*, including median/interquartile boxplots and jittered sample points.

### Single-Cell RNA-seq Analysis of EGFR-Mutant UCSF Cohort

Single-cell RNA-seq data from the internal UCSF EGFR-mutant cohort were processed using Python (v3.8+) and Scanpy. Cells were stratified by patient group: *CDK4*amp (n = 3) vs. *CDK4/6*wt (n = 11) cases. Cells annotated as cancer via inferCNV-based labeling were retained for downstream analyses. UMAP coordinates were generated using single cells from *CDK4*amp cases using a reproducible pipeline implemented in Python: normalization (sc.pp.normalize_total), log-transformation, PCA, neighborhood graph construction, and UMAP embedding.

Expression of genes of interest (GOI: *AGR2*, *ASNS*, *CCZ1B*, *CDK4*, *GGCT*, *PSMA2*, and *STEAP1*) was visualized on UMAPs using scaled expression values.

Log-normalized expression of GOIs in cancer cells was compared between *CDK4*amp and *CDK4/6*wt groups. Violin and strip plots were generated using Seaborn in Python. Medians and interquartile ranges were annotated. Between-group comparisons were assessed using Mann–Whitney U tests.

For each group, Spearman correlation coefficients (ρ) between *CDK4* and other GOIs were calculated using Python’s SciPy library, with two-sided significance assessed using Spearman’s rank test. Differences in correlation coefficients between *CDK4*amp and *CDK4/6*wt groups (βρ) were assessed by comparing Fisher z-transformed correlation coefficients using a z-test. P-values from between-group comparisons were adjusted for multiple testing using the Benjamini–Hochberg false discovery rate (FDR) method.

An EMT score was computed per cell as the mean expression of canonical EMT markers (*VIM*, *ZEB1*, *SNAI1*, *TWIST1*, *CDH2*, *FN1*). Correlations between EMT score and mean GOI expression were analyzed and visualized by group using violin and scatter plots with regression fits generated in Python. Group-specific Spearman ρ and p-values were annotated to assess differential EMT–GOI associations.

Scatter plots were generated for each GOI to assess EMT score correlations separately in *CDK4*amp and *CDK4/6*wt cancer cells. Each plot displays the Spearman ρ and p-value to illustrate gene-specific EMT associations across patient groups.

### Statistical analyses

We used One-way ANOVA with Tukey’s multiple comparisons and Student’s T-test to assess statistical significance in preclinical functional studies (GraphPad Prism); variations across samples were expressed as S.E.M. For the Fraction of Genome Alterations (FGA) and Tumor Mutational Burden (TMB) analyses, the Wilcoxon non-parametric two-group comparison test and two-sided Student t-test assessed statistical significance. Statistics were computed using Fisher’s exact test for the mutual exclusivity and WGD analyses. For the pathway enrichment analysis of SNVs, statistics were calculated using a hypergeometric test (phyper in R).

### Data Repositories

DNA sequencing datasets included in the manuscript are available at the NCBI SRA reference accession code PRJNA1093094 (release date April 28th, 2025). RNA sequencing datasets in the manuscript are available at the NCBI GEO reference accession code: GSE262270 (release date TBD).

## Acknowledgments

This project was supported by the NIH/NCI U54CA224081 (T.G.B), The Damon Runyon Cancer Research Foundation P0528804 (United States), Doris Duke Charitable Foundation P2018110 (United States), V. Foundation P0530519 (to C.M.B.). The authors thank Thomas B.K. Watkins for critical advice in the whole genome doubling analysis, Foundation Medicine Inc. (Cambridge, Massachusetts) for supporting genomic data, Kari Herrington from the UCSF Microscopy Core for the helpful advice with the immunofluorescence experiments, Robin Lea for the helpful advice with the clinical dataset analyses. We gratefully acknowledge the use of data from the MSK-IMPACT GENIE BioPortal, which provided valuable genomic datasets essential for this study.

## Conflicts of Interest

C.M.B receives research funding from AstraZeneca, Novartis, Roche, and Puma.

## Notes

### Summary of Updates

Clarified methods in figure legends, added to discussion section, fixed typographical errors.

