## Supplementary Figures for "CDK4 or CDK6 upregulation induces DNA replication stress and genomic instability to cause EGFR targeted therapy resistance in lung cancer"

Supplementary Figure 1

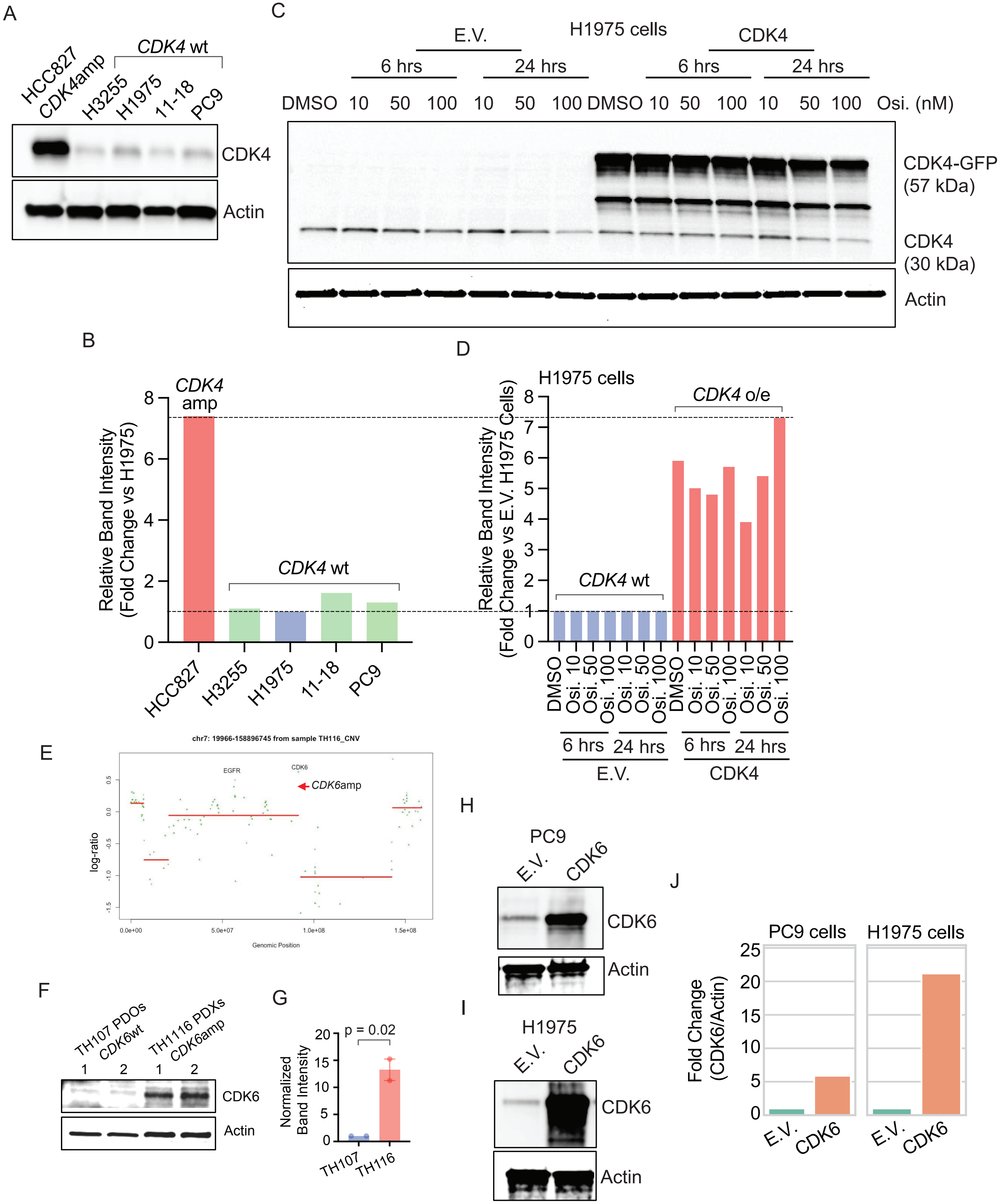

**Supplementary Figure 1**

(A, B) Immunoblotting probing for CDK4 protein (A) and relative quantification normalized to Actin and expressed as a fold change compared to H1975 CDK4 expression (B), in *EGFR* mutant LUAD cells carrying *CDK4*amp (HCC827) versus *CDK4* wild-type cells (H3255, H1975, 11-18, PC9). (C, D) Immunoblotting probing for CDK4 protein (C) and relative quantification normalized to Actin and expressed as a fold change compared to H1975 empty vector control (E.V.) (D) comparing H1975 cells overexpressing CDK4 versus cells carrying empty vector control and treated with DMSO or osimertinib at the indicated dose. (E) Whole exome sequencing data demonstrating *CDK6* amplification in the TH116 (EGFR p.L858R) patient derived xenograft (PDX). (F) Immunoblotting demonstrating CDK6 protein expression in TH116 PDXs with relative quantification (G) normalized to Actin and expressed as a fold change compared to TH107 (EGFR exon 19 del, CDK4/6wt) patient-derived organoids (PDOs) (Student's t-test; error bars representing SEM). (H-J) Immunoblotting demonstrating CDK6 protein expressing, in CDK6 overexpressing PC9 (H) and H1975 (I) cells compared to cells carrying empty vector control (E.V.) with quantification normalized to Actin expression (J).

Supplementary Figure 2

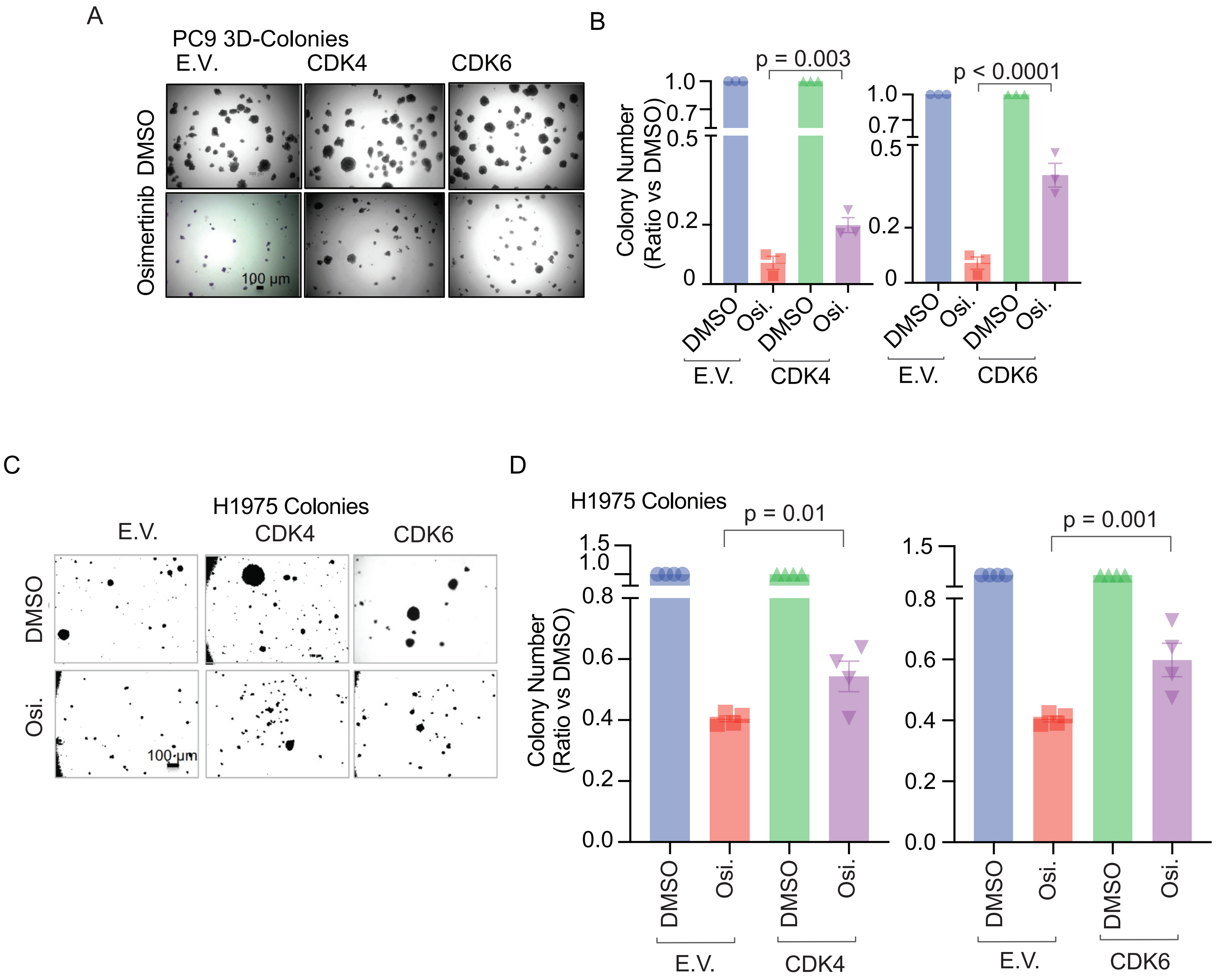

**Supplementary Figure 2**

Soft agar colony formation assay and quantification using CDK4/6 overexpressing PC9 (A, B), and H1975 (C, D) EGFRmt LUAD cell lines and E.V. control cells (p-value calculated with one-way ANOVA and Tukey's multiple comparisons test; error bars representing SEM). (E.V.: empty vector; CDK4 and CDK6: CDK4 or CDK6 overexpressing cells; Osi.: osimertinib; for the 3D-soft agar colony formation assay, osimertinib was used at 50 nM for PC9 and H1975 cells).

Supplementary Figure 3

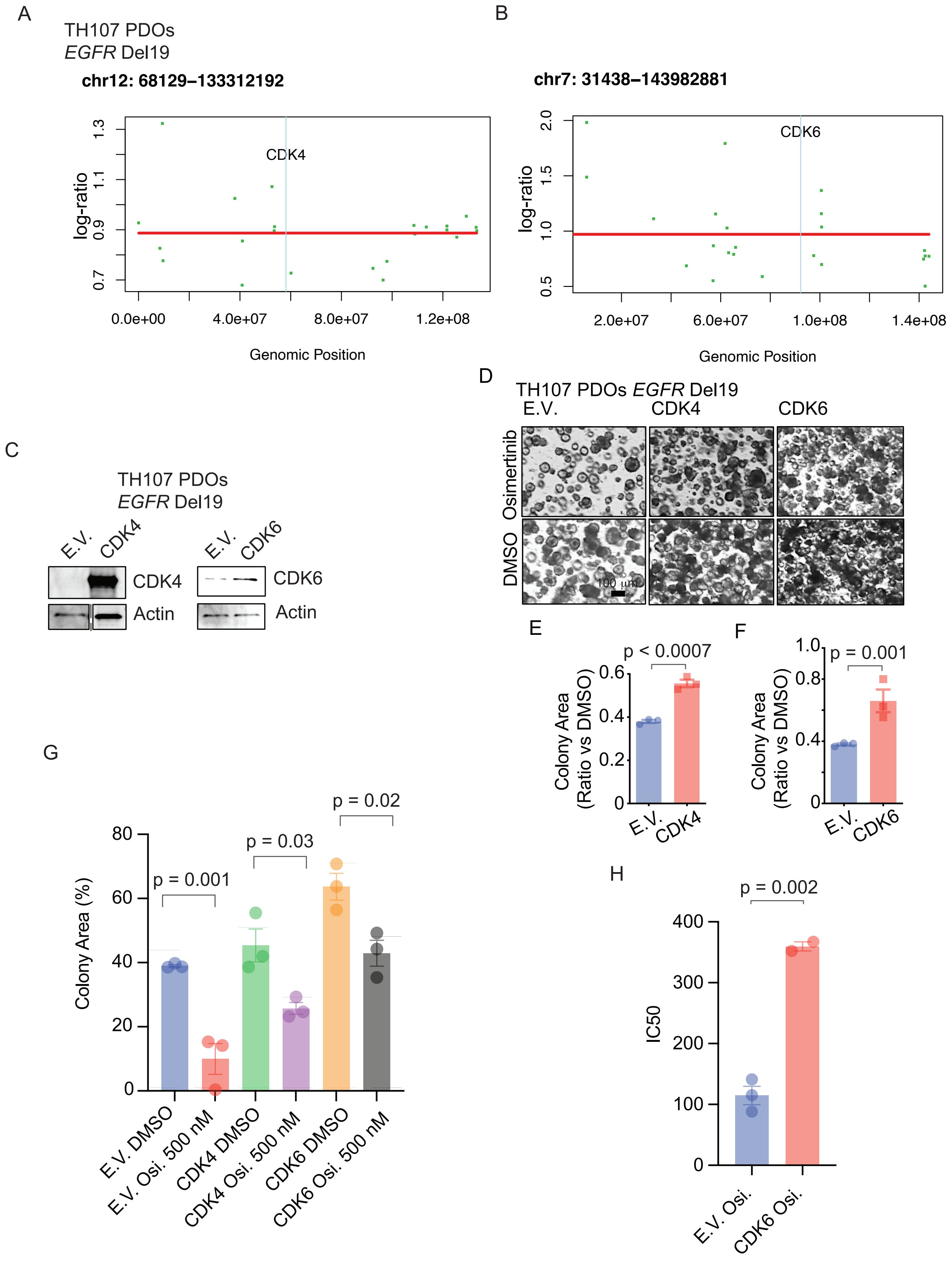

**Supplementary Figure 3**

Assessment of *CDK4* (A) and *CDK6* (B) copy number status in TH107 PDOs (*CDK4* and *CDK6* copy number computed aligning whole exome sequencing reads, represented as dots, to the corresponding gene's locations on chromosomes 12 and 7; the absence of *CDK4/6* copy number gains is highlighted with a gray bar corresponding to the *CDK4/6* gene locations). (C) Immunoblot analysis demonstrating CDK4 and CDK6 overexpression in genetically modified *EGFR* Del19 TH107 PDOs. (D-G) Representative images (D) and colony area quantification (E-G) upon long-term (35 days) osimertinib (500 nM) treatment with CDK4/6 overexpressing, *EGFR* Del19 TH107 PDOs, compared to DMSO treated controls (error bars representing SEM). (H) IC50 with osimertinib treatment in *EGFR* Del19, E.V., or CDK6 overexpressing TH107 PDOs (Student's t-test; error bars representing SEM). (E.V.: empty vector; CDK4 and CDK6: CDK4 or CDK6 overexpressing PDOs; Osi.: osimertinib).

Supplementary Figure 4

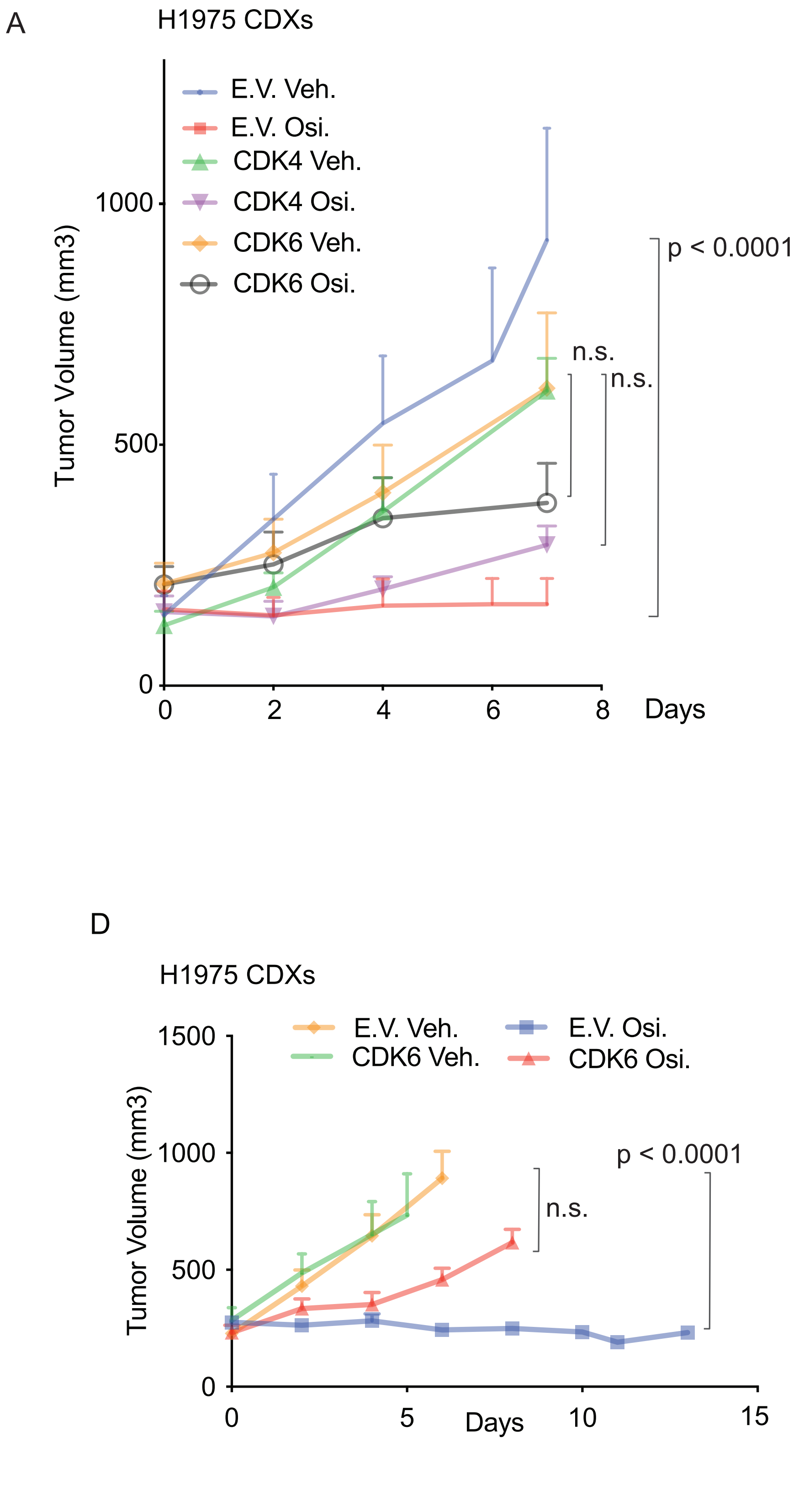

**Supplementary Figure 4**

(A-C) Growth curves (A) and percentage of tumor volume change (B, C) of 7 day osimertinib (5 mg/kg) or vehicle (Veh.) treated CDK4 (A, B) or CDK6 (A, C) overexpressing H1975 CDXs compared to empty vector (E.V.) controls (n = 4-6 xenografts per group; error bars representing SEM). (D,E) Growth curves (D) and percentage of tumor volume change (E) of 14 day osimertinib (5 mg/kg) or vehicle (Veh.) treated CDK6 overexpressing H1975 CDXs compared to empty vector (E.V.) controls (n = 4-6 xenografts per group; error bars representing SEM)
(E.V.: empty vector; CDK4: CDK4 overexpressing CDXs; CDK6: CDK6 overexpressing CDXs; Veh.: vehicle; Osi.: osimertinib).

Supplementary Figure 5

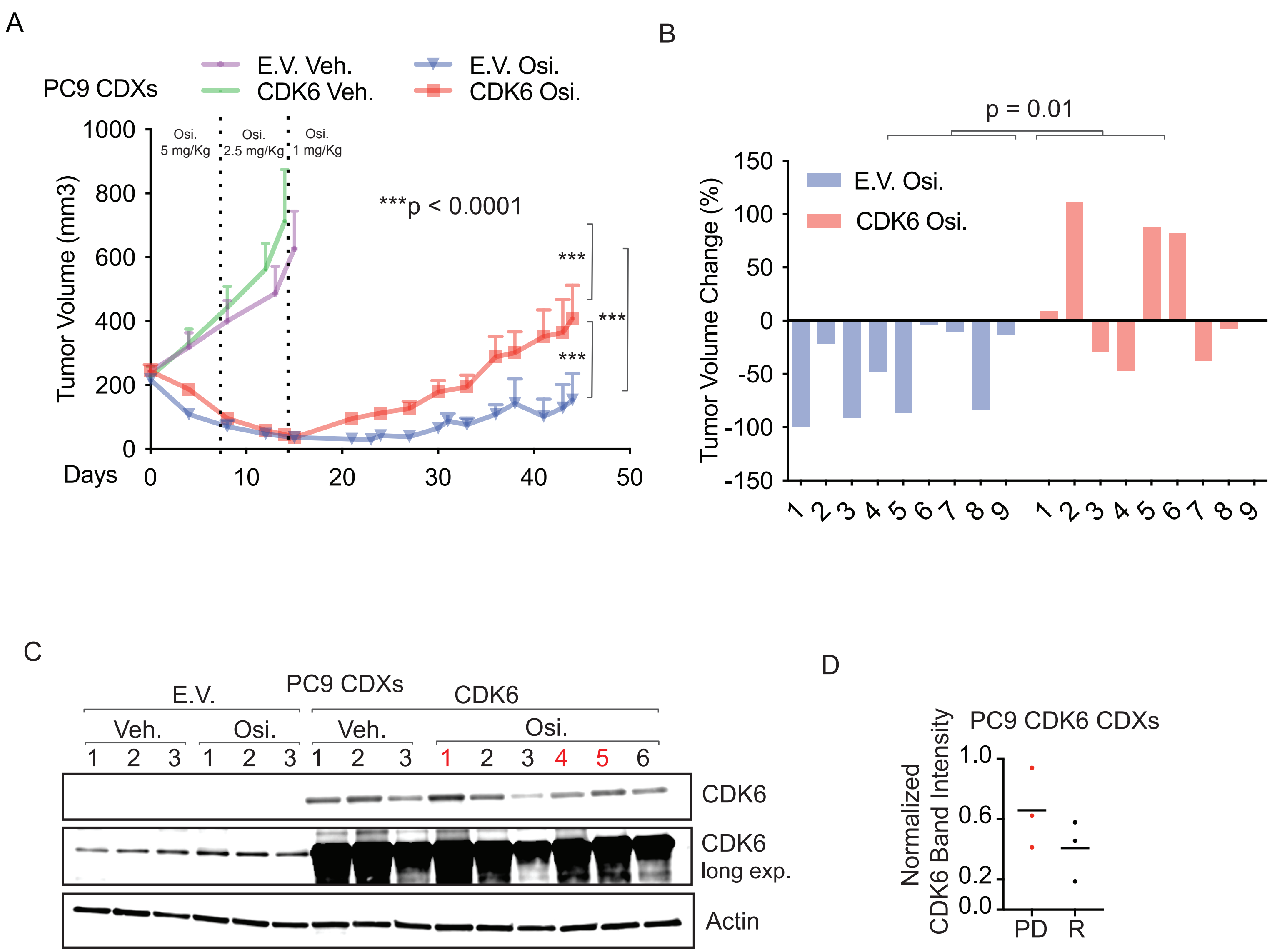

**Supplementary Figure 5**

(A,B) Growth curves (A) and individual xenograft response (B) of long-term, osimertinib-treated (1 mg/kg to 5 mg/kg) PC9 CDXs, overexpressing CDK6 compared to empty vector controls (n = 5 xenografts per vehicle groups and n = 5-10 xenografts per osimertinib-treated groups; one-way ANOVA with Tukey's multiple comparisons test and Student's t-test; error bars representing SEM). (C, D) Immunoblot analysis of CDK6 protein expression in progressive disease (PD) and responsive (R) CDK6 overexpressing PC9 CDXs versus E.V. control (C) and quantification (D) (dot plots show average values, red label/dot indicates a progressive disease (PD) CDX.

(E.V.: Empty Vector; CDK4: CDK4 overexpressing CDXs; CDK6: CDK6 overexpressing CDXs; Osi.: osimertinib).

Supplementary Figure 6

A

H1975 CDXs

E2F Targets

B

G2/M Checkpoint

C

MYC TARGETS V1

D

DNA Repair

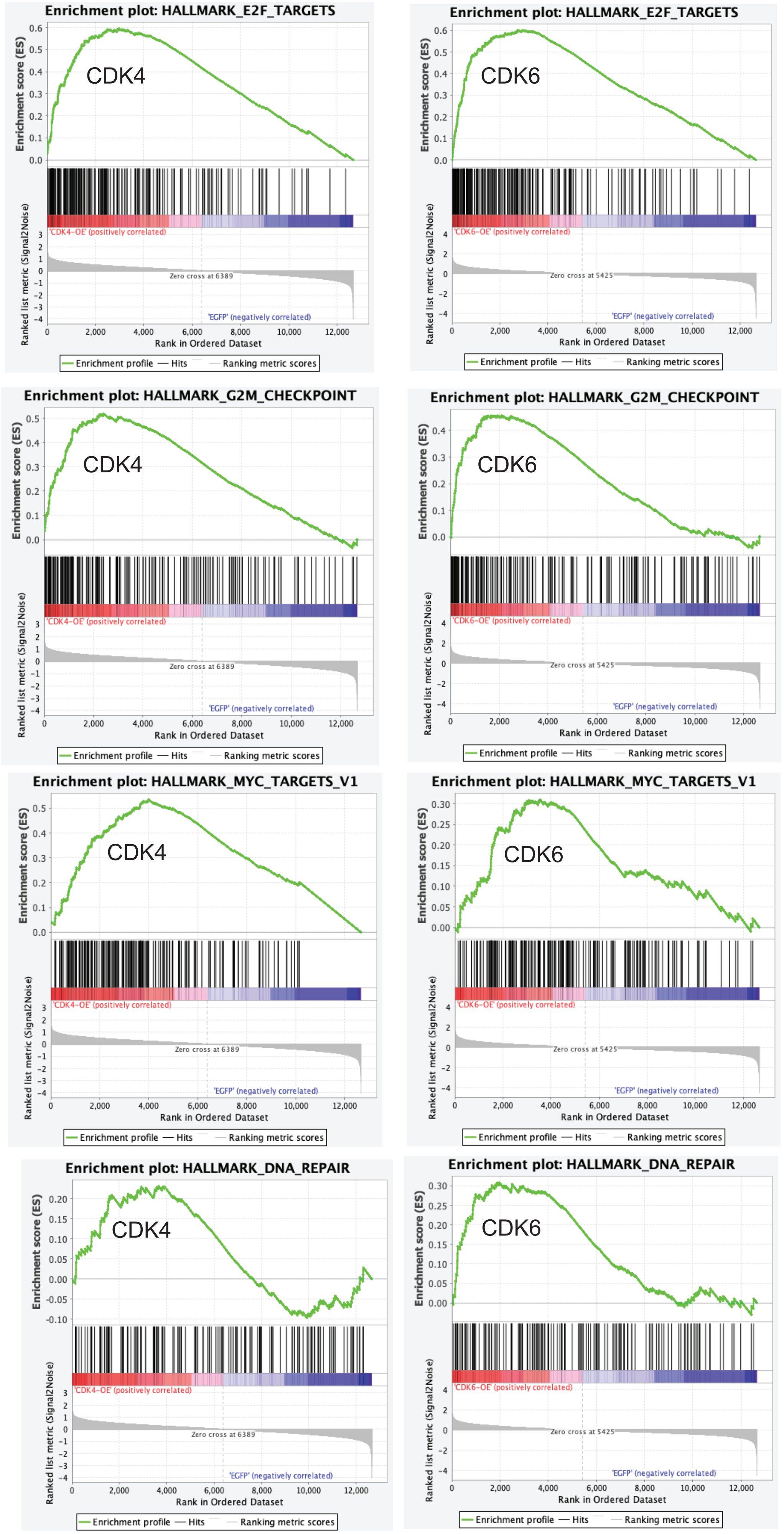

**Supplementary Figure 6**

(A-D) Gene Set Enrichment Analysis (GSEA) of H1975 CDK4/6 overexpressing CDXs compared to empty vector (E.V.) controls; enrichment in E2F targets (A), G2/M checkpoint (B), MYC TARGETS V1 (C), and DNA repair (D) pathways were observed. (E.V.: empty vector; CDK4: CDK4 overexpressing CDXs; CDK6: CDK6 overexpressing CDXs).

Supplementary Figure 7

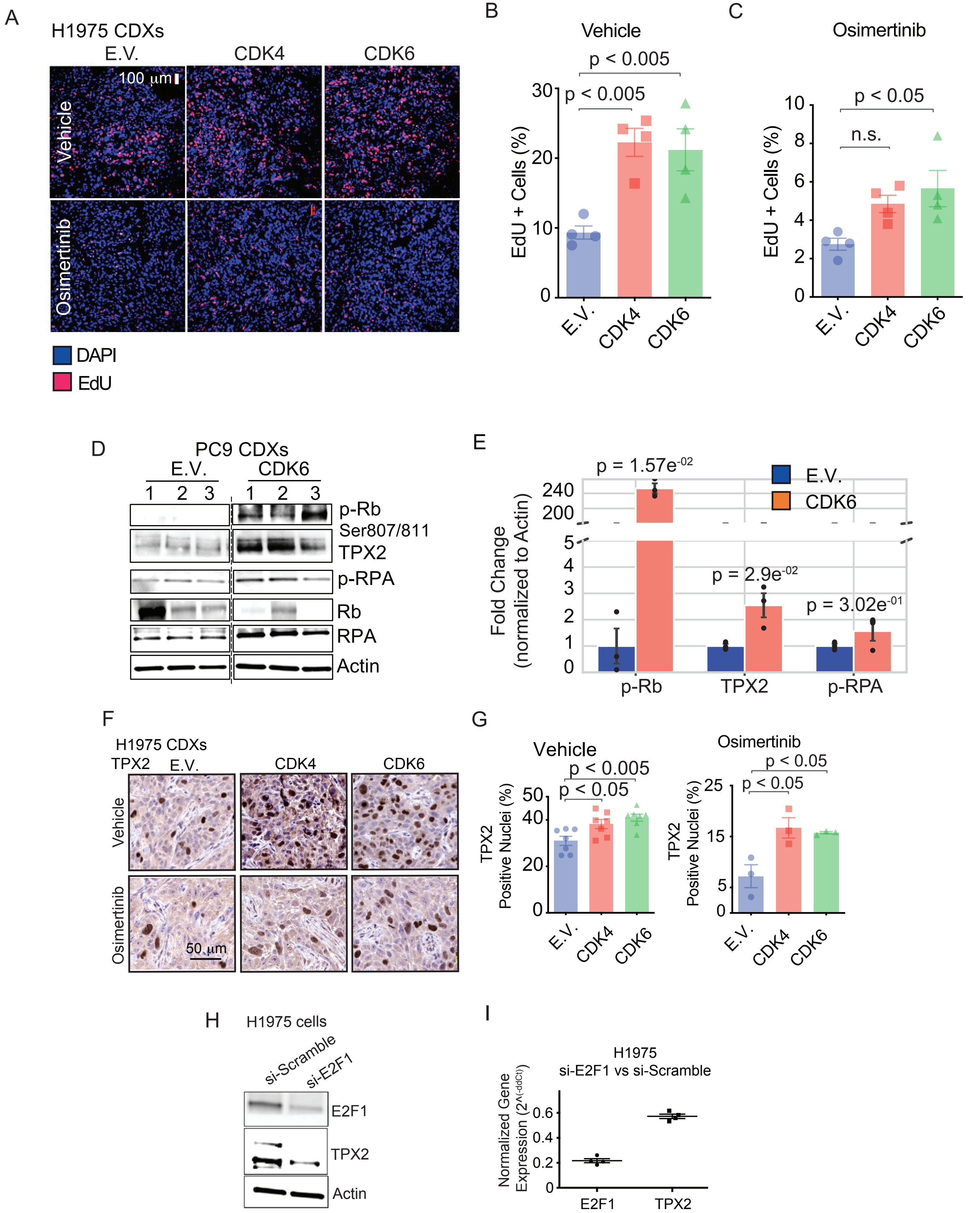

**Supplementary Figure 7**

(A-C) S-phase and active cell cycle progression analysis in H1975 CDK4/6overexpressing tumors by quantification of EdU incorporation (A, red dots) in newly synthesized DNA in the vehicle (B) and osimertinib-treated (C) tissues (n = 4 images for each xenograft per group; p-value assessed by one-way ANOVA and Tukey's multiple comparisons tests; error bars representing SEM). (D) Immunoblot analysis of replication stress biomarkers in CDK6 overexpressing PC9 CDXs versus E.V. controls (D) and protein expression quantification (E) (normalized to Actin and expressed as a fold change vs. E.V.; error bars representing SEM; each black dot represents a replicate; Welch's t-test log transformed). (F) Representative images of TPX2 IHC stained H1975 CDXs and corresponding quantification (G) of vehicle and osimertinib treated tumors comparing CDK4/6 overexpressing to E.V. controls (n = 3-7 xenografts per group; p value calculated with one-way ANOVA and Tukey'smultiple comparisons test; error bars representing SEM). (H, I) E2F1 knockdown using siRNA in H1975 cells (H) and normalized gene expression assessed by qPCR for E2F1 and TPX2 mRNAs in si-E2F1 cells versus si-scramble (I) (error bars representing SEM). (E.V.: Empty Vector; CDK4: CDK4 overexpressing CDXs; CDK6: CDK6 overexpressing CDXs or cells).

Supplementary Figure 8

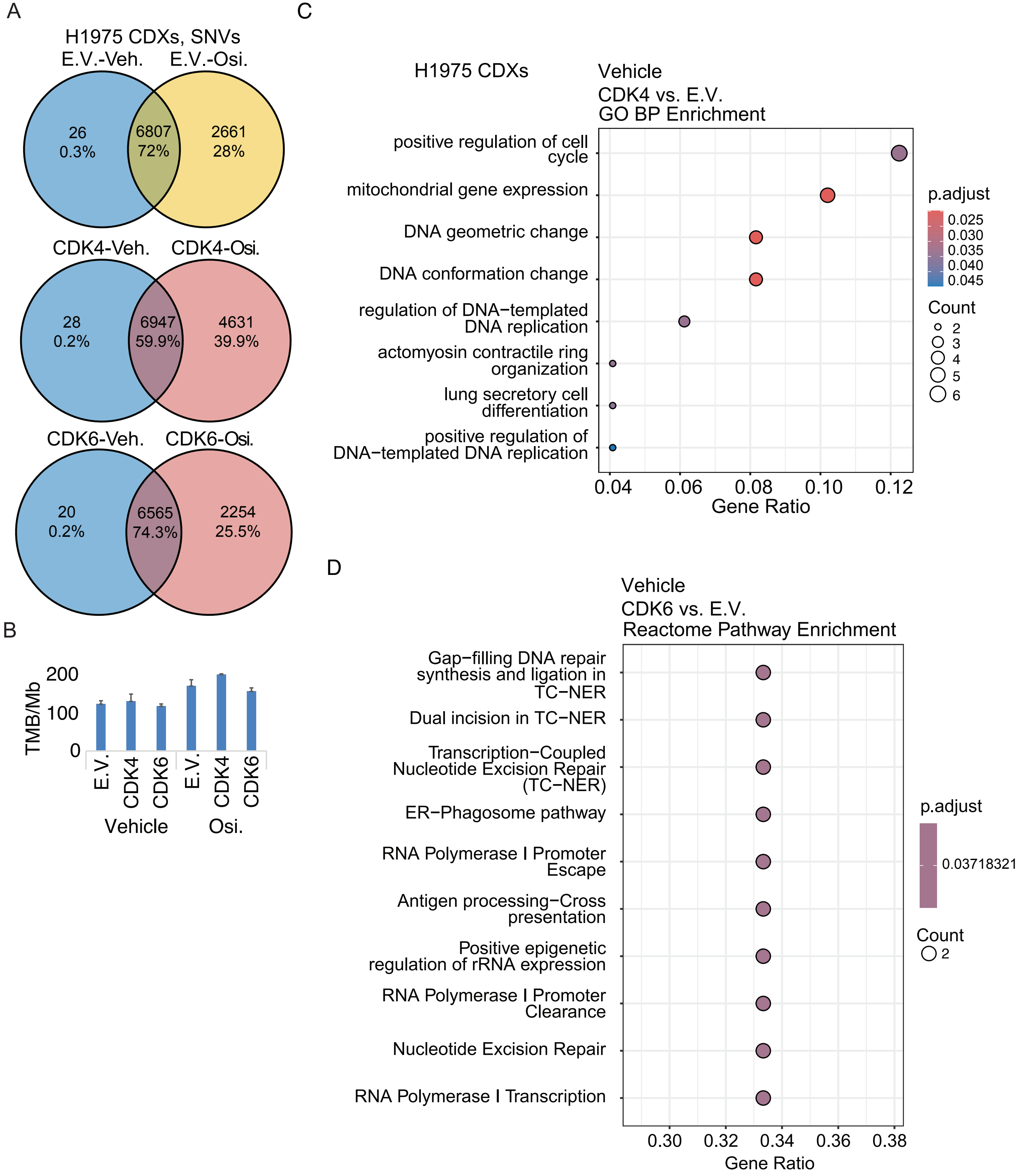

**Supplementary Figure 8**

(A, B) Venn diagrams (A) representing the total number with percentages of unique and shared SNVs detected in E.V., CDK4, and CDK6 overexpressing CDXs treated with osimertinib or vehicle control; no differences in TMB (B) were observed comparing E.V. with CDK4 and CDK6 overexpressing CDXs in the corresponding treatment group (n = 2 xenografts for each group). (C, D) Gene ontology (GO) biological process, and Reactome pathway enrichment analyses were performed on the set of overlapping genes identified from CDK4 (C) or CDK6 (D) upregulated and copy number-altered datasets. The studies revealed significant enrichment in pathways related to cell cycle regulation, DNA replication, and repair. Dotplots illustrate the top 10 enriched categories for each database, highlighting key biological processes and signaling pathways implicated in CDK4/6-driven tumor biology (n = 2 xenografts for each group for the WES/CNAs analysis; n = 3 xenografts for each group for the RNAseq analysis; data normalized to E.V. controls).
(E.V.: empty vector; CDK4 and CDK6: CDK4 or CDK6 overexpressing CDXs; Veh.: vehicle; Osi.: osimertinib 5 mg/kg).

Supplementary Figure 9

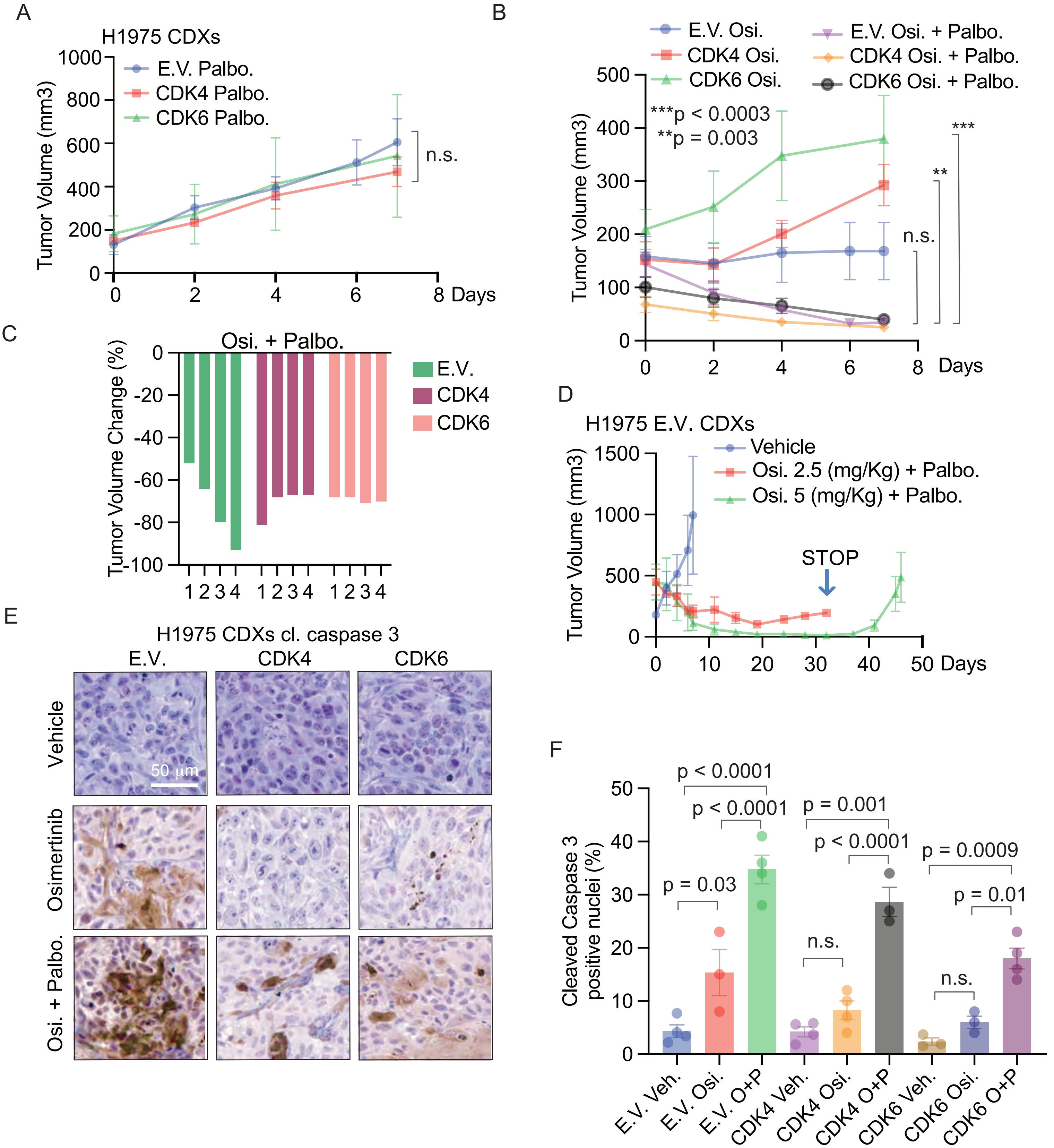

**Supplementary Figure 9**

(A) Growth curves of E.V. and CDK4/6 overexpressing H1975 CDXs, treated with palbociclib (100 mg/kg) (error bars representing SEM). (B, C) Growth curves (B) and individual tumor volume changes (C) of CDK4/6 or empty vector H1975 CDXs treated with osimertinib (5 mg/kg) or osimertinib (5 mg/kg) in combination with palbociclib (100 mg/kg) (n = 3-4 xenografts per group; error bars representing SEM). (D) Growth curves of E.V. H1975 CDXs treated with palbociclib in combination with osimertinib (Palbo.: palbociclib 150 mg/kg; Osi.2.5/Osi.5: osimertinib 2.5-5 mg/kg; n = 2 xenografts per group). (E, F) Representative images (E) and IHC staining quantification (F) of cleaved caspase 3 from CDK4/6 overexpressing H1975 CDXs treated with osimertinib (5 mg/kg) and combinatorial treatments with palbociclib (100 mg/kg) compared to empty vector controls (n = 3-4 xenografts per group; error bars representing SEM). (E.V.: empty vector; CDK4 and CDK6: CDK4 or CDK6 overexpressing CDXs; Osi.: osimertinib; Palbo.: palbociclib; p-value determined using one-way ANOVA with Tukey's multiple comparisons test).

Supplementary Figure 10

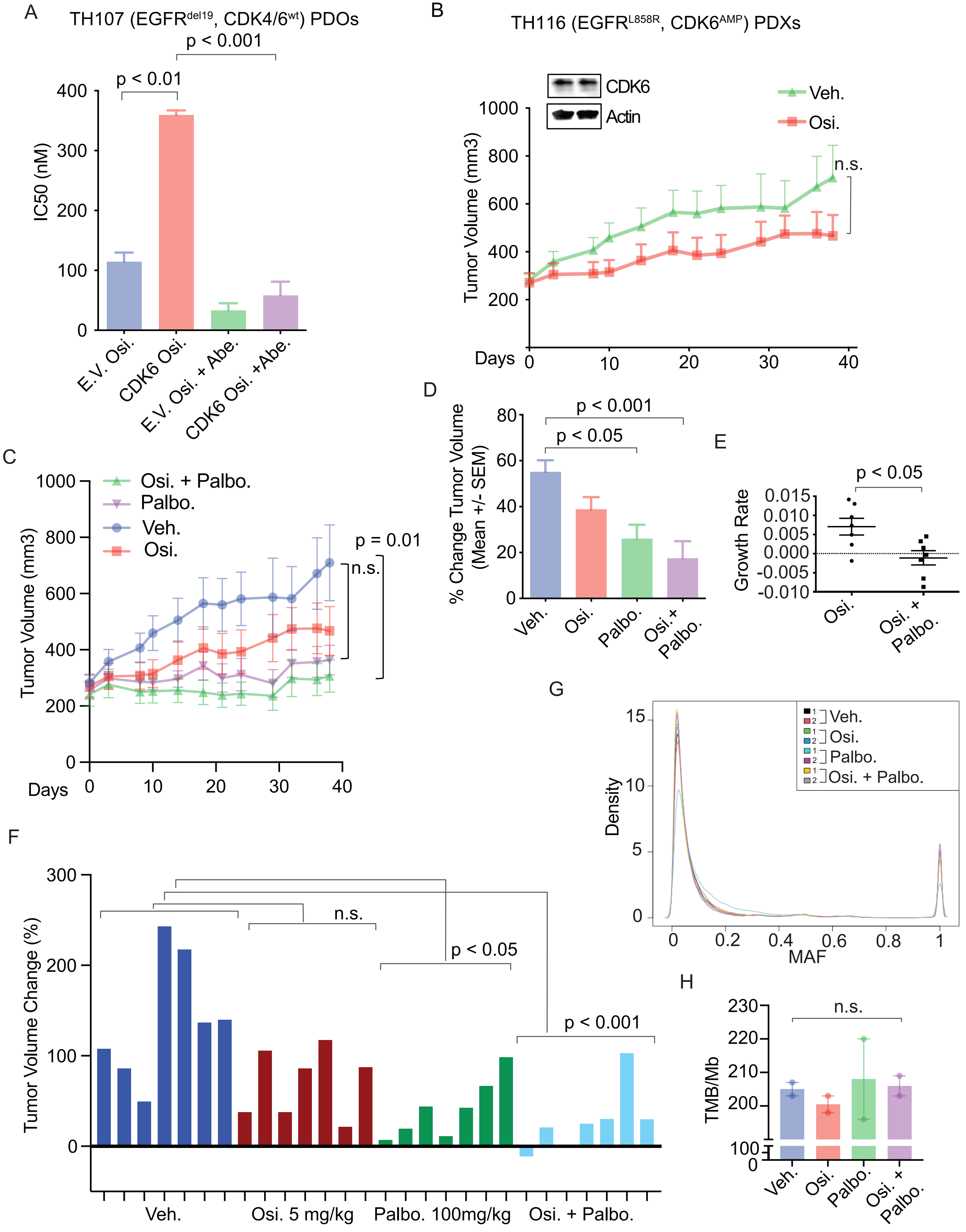

**Supplementary Figure 10**

(A) IC50 with mono- and combination treatment of osimertinib and the CDK4/6 inhibitor abemaciclib (1 µM) in *EGFR* Del19, E.V. or CDK6 overexpressing TH107 PDOs (error bars representing SEM). (B) Growth curves for TH116 Patient-Derived Xenografts (PDXs), harboring EGFR p.L858R and *CDK6* amplification, treated with vehicle or osimertinib (5 mg/kg) (n= 7 xenografts per group; Student's t-test). (C-E) Growth curves (C), percentage of tumor volume change (D), and growth rate (E) of TH116 PDXs treated with osimertinib (5 mg/kg) or a combination of osimertinib (5 mg/kg) and palbociclib (100 mg/Kg, Osi. + Palbo.) (Student's t-test). (F) Percentage of tumor volume change of TH116 PDXs treated with vehicle, osimertinib (5 mg/kg), palbociclib (100 mg/kg), or combination of osimertinib and palbociclib (Osi. + Palbo.)(n= 7 xenografts per group). (G, H) Mutational Allele Frequency (G) and TMB (H) analyses in TH116 PDXs, treated with vehicle, osimertinib (5 mg/kg), palbociclib (100 mg/kg), or a combination of osimertinib and palbociclib (Osi. + Palbo.) (2 xenografts per group). (E.V.: empty vector; CDK4 and CDK6: CDK4 or CDK6 overexpressing CDXs or PDOs; Osi.: osimertinib; Abe: abemaciclib; p-value determined using one-way ANOVA with Tukey's multiple comparisons test; error bars representing SEM).

Supplementary Figure 11

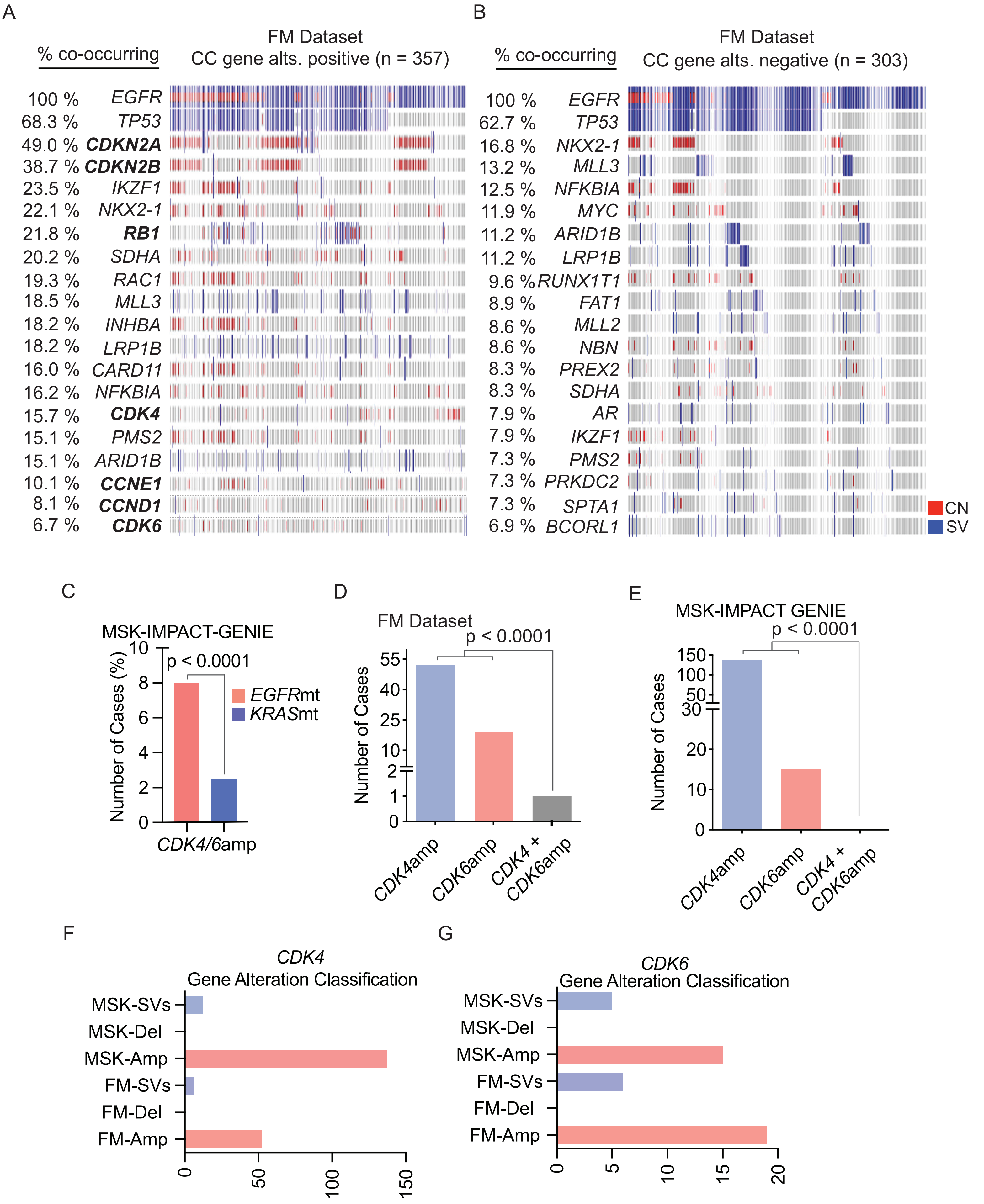

**Supplementary Figure 11**

(A, B) Oncoprints highlighting recurrent gene alterations detected in EGFRmt LUADs harboring cell cycle (CC) gene alterations (CC pos.; A) or negative for such alterations (CC neg.; B) (FM, Foundation Medicine dataset). (C) Frequency of concurrent *CDK4/6* gene alterations in *EGFR* mutant and *KRAS* mutant LUAD cases (MSK-Impact GENIE Dataset). (D, E) Mutual exclusivity analysis of *CDK4* and *CDK6* gene alterations in EGFRmt tumors from Foundation Medicine (FM) (D) or MSK-Impact GENIE (E) targeted exome sequencing (p-value calculated with Fisher's test). (F, G) Classification of alterations affecting *CDK4* (F) and *CDK6* (G) genes in Foundation Medicine (FM) or MSK-Impact GENIE targeted exome sequencing datasets. Somatic variant (SV), deletion (Del), amplification (Amp).

Supplementary Figure 12

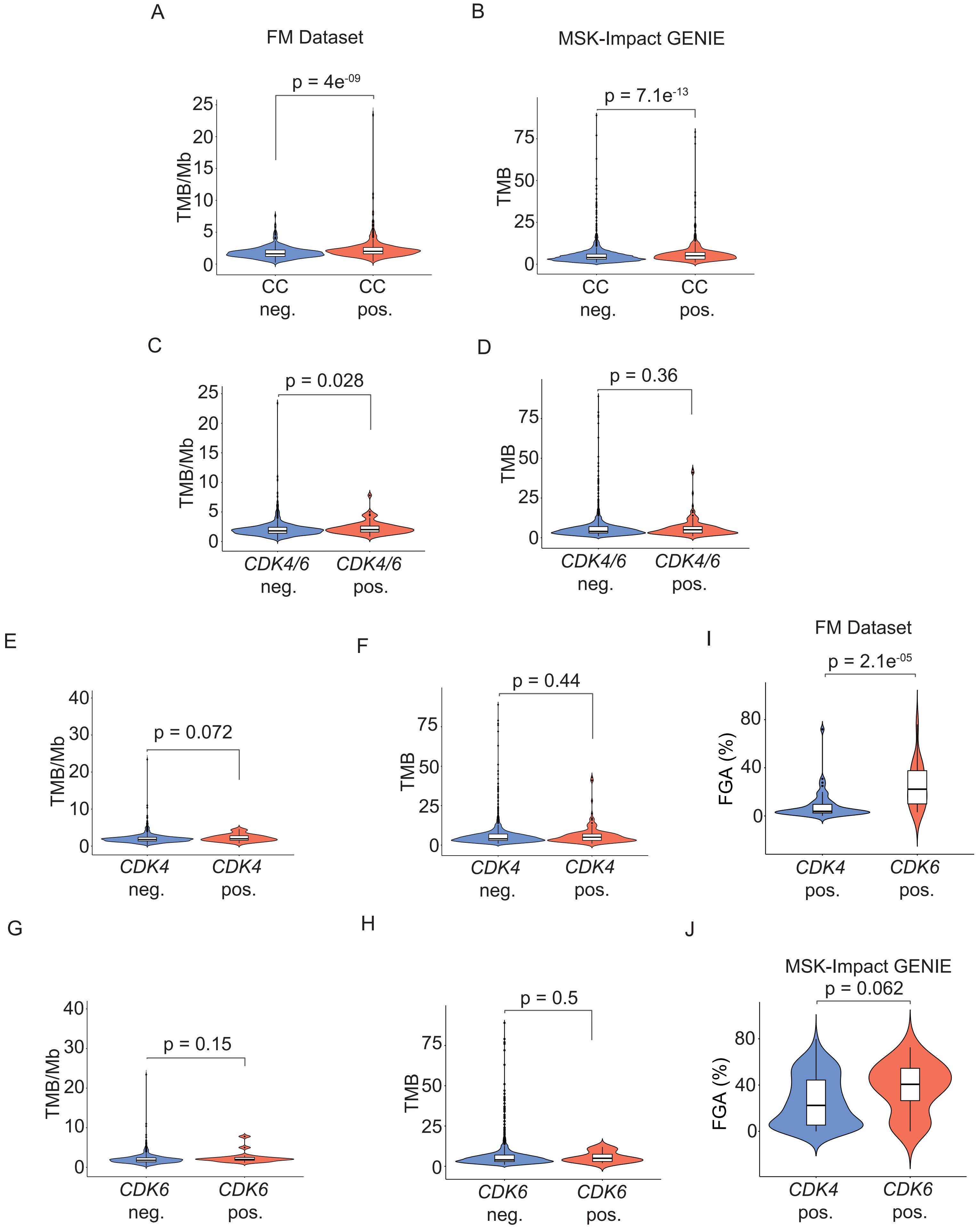

**Supplementary Figure 12**

(A-H) TMB analyses in EGFRmt LUAD using Foundation Medicine (FM) targeted exome sequencing (A, C, E, G) or MSK-Impact GENIE (B, D, F, H) datasets, stratifying tumors carrying concurrent cell cycle CNAs (CC pos., A, B), *CDK4/6*amp (*CDK4/6* pos., C, D), *CDK4*amp (*CDK4/6*amp pos., E, F) or *CDK6*amp (*CDK6*amp pos., G, H) versus tumors negative for such alterations (CC/*CDK4/6*amp/*CDK4*amp/ *CDK6*amp neg.). (I, J) FGA analysis in EGFRmt tumors using Foundation Medicine (FM) (I) or MSK-Impact-GENIE (J) targeted exome sequencing datasets, comparing tumors carrying *CDK4* or *CDK6* CNAs (*CDK4* or *CDK6* pos.).

(TMB and FGA boxplots included in violin plots represent the interquartile range, lower quartile 25%, median, and upper quartile 75%; p-value calculated with Wilcoxon test).

See also Supplementary Table 3.

Supplementary Figure 13

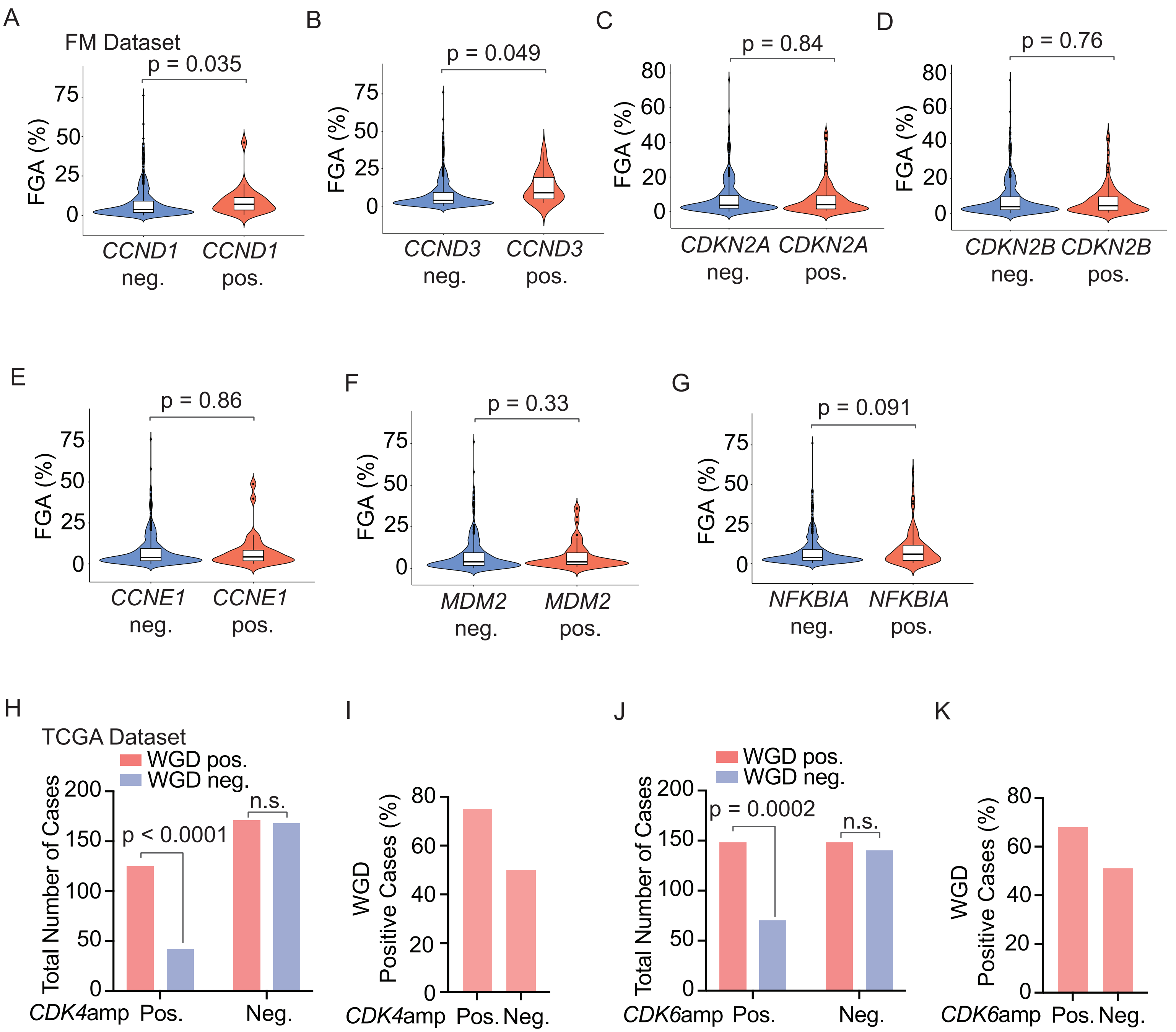

**Supplementary Figure 13**

(A-G) FGA analysis with Foundation Medicine dataset comparing *CCND1* (A), *CCND3* (B), *CDKN2A* (C), *CDKN2B* (D), *CCNE1* (E), *MDM2* (F) or *NFKIB* (G) CNAs positive versus negative EGFRmt LUAD cases (n = 63 for *MDM2* pos. CNAs; n = 94 for *NFKIB* pos. CNAs; n = 660 total EGFRmt cases; see Supplementary Table 3 for the number of cases per cell cycle gene alteration). (H-K) Whole Genome Doubling (WGD) analysis using TCGA data from NSCLC cases carrying *CDK4* (H, I) or *CDK6* (J, K) CNAs or negative for such alterations (p-value calculated with Fisher's test).

See also Supplementary Table 3.

Supplementary Figure 14

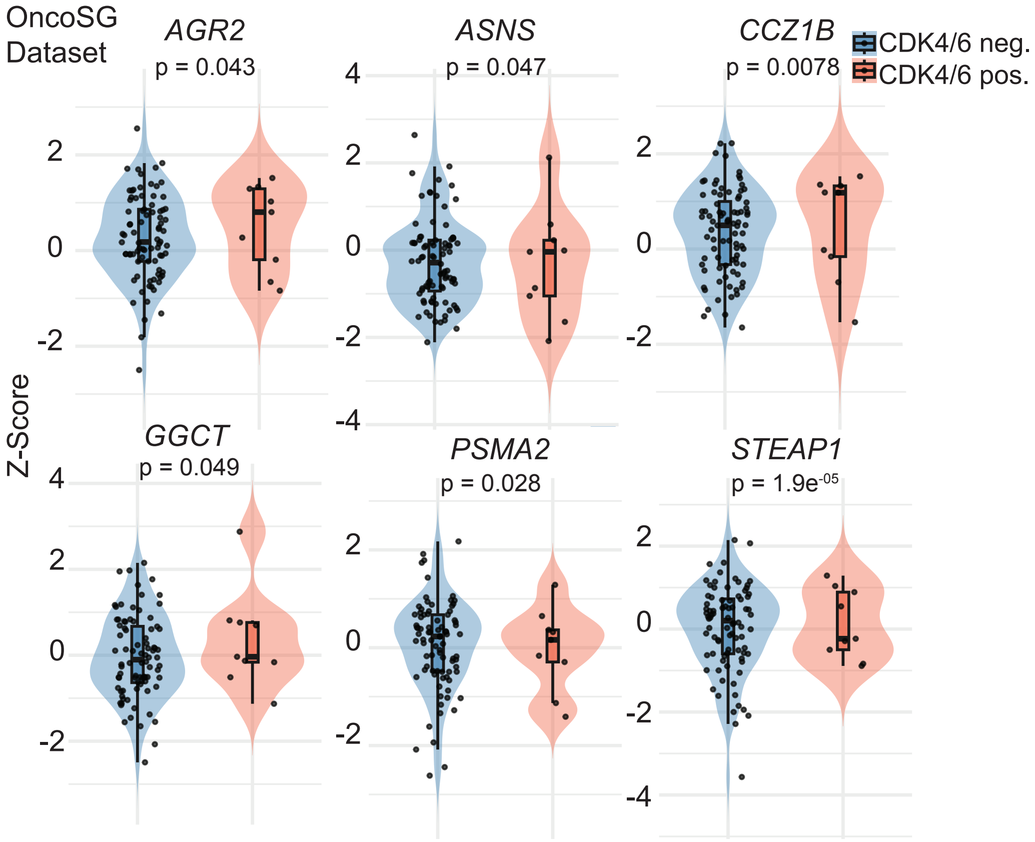

**Supplementary Figure 14**

Violin plots display Z-score–normalized gene expression values for six selected genes: *AGR2*, *ASNS*, *CCZ1B*, *GGCT*, *PSMA2*, and *STEAP1*, across OncoSG EGFRmt LUAD cases, CDK4/6 positive (n = 9) or negative (n = 84) subgroups. Each violin plot shows the distribution of expression values, overlaid with boxplots (showing medians and interquartile ranges) and individual sample-level data (associated p-value from a two-sample unpaired Welch's t-test).

Supplementary Figure 15

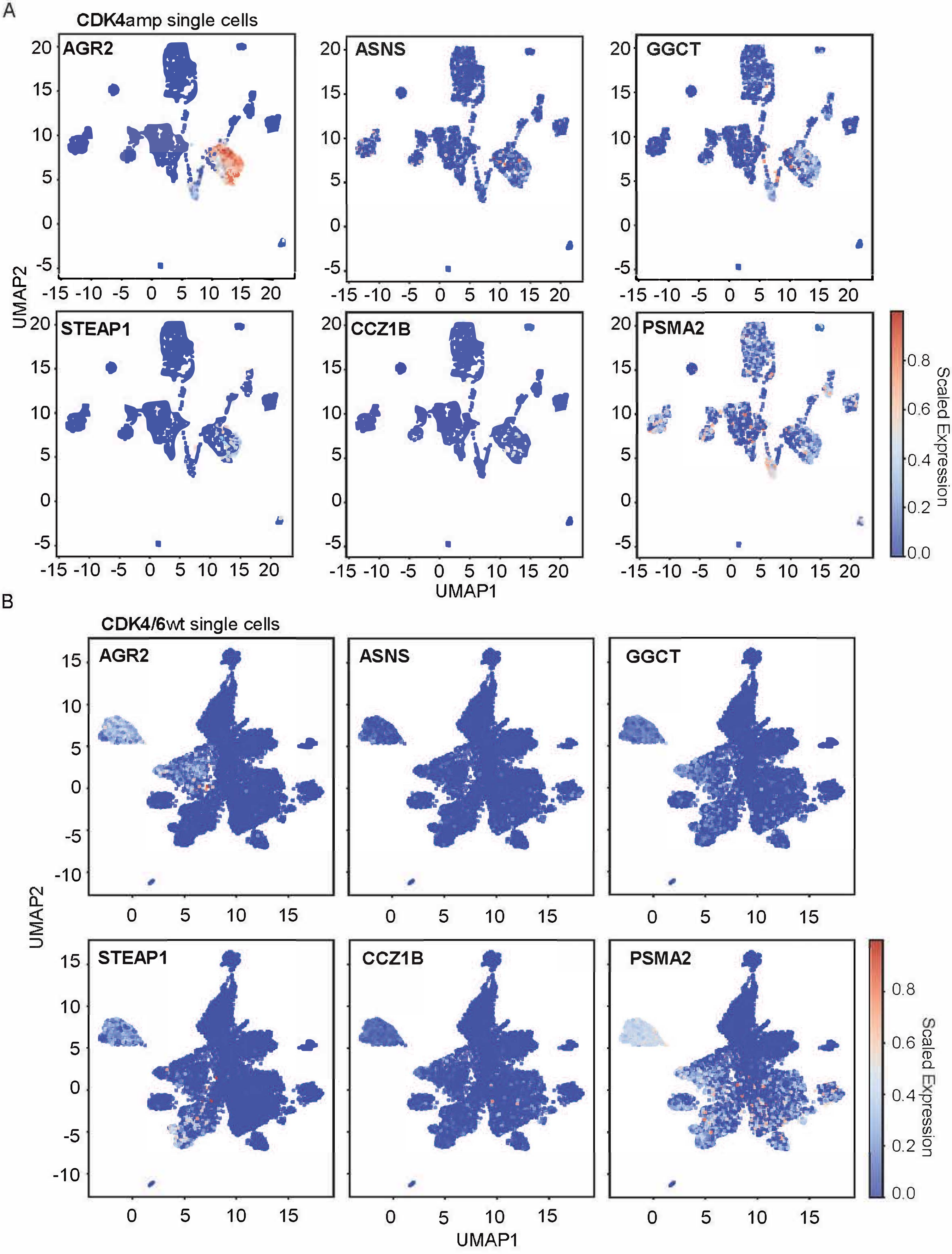

**Supplementary Figure 15**

(A, B) UMAP plots displaying the spatial distribution of *AGR2*, *ASNS*, *GGCT*, *STEAP1*, *CCZ1B*, and *PSMA2* expression within *CDK4*amp (A) and *CDK4/6*wt (B) clinical cases' single cells, highlighting upregulation in the *CDK4*amp group.
