## Supplementary Tables for "CDK4 or CDK6 upregulation induces DNA replication stress and genomic instability to cause EGFR targeted therapy resistance in lung cancer"

Supplementary Table 1 – Cell cycle gene alterations detected in parental EGFRmt NSCLC preclinical models

| **NSCLC model** | **EGFR mutation(s)** | **Cell Cycle Gene Alterations** |
|---|---|---|
| H1975 | p.EGFR L858R, T790M | *CDKN2A* SNV p.E69Ter |
| PC9 | *EGFR* exon 19 del (E756-A750) | *CDKN2A* SNV p.G67V |
| HCC827 | *EGFR* exon 19 del (E756-A750) | *CDKN2C* CN gain<br>*CDK4* CN gain |
| TH116 | p.EGFR L858R | *CCND1* SNV p.Y226F<br>*CDK6* CN gain |
| TH107 | *EGFR* exon 19 del (E746_A750>QP) | *CCND3* SNV p.S187A |

Supplementary Table 2 – EGFRmt NSCLC preclinical models

| *In vitro* 3-D models | Oncogenic Driver | Treatment | |
|---|---|---|---|
| 1 H1975 E.V. | *EGFR* L858R, T790M | DMSO | Osimertinib |
| 2 H1975 CDK4 | *EGFR* L858R, T790M | DMSO | Osimertinib |
| 3 H1975 CDK6 | *EGFR* L858R, T790M | DMSO | Osimertinib |
| 4 HCC827 E.V. | *EGFR* Del19 | DMSO | Osimertinib |
| 5 PC9 E.V. | *EGFR* Del19 | DMSO | Osimertinib |
| 6 PC9 CDK4 | *EGFR* Del19 | DMSO | Osimertinib |
| 7 PC9 CDK6 | *EGFR* Del19 | DMSO | Osimertinib |
| 8 TH107 E.V. | *EGFR* Del19 | DMSO | Osimertinib |
| 9 TH107 CDK4 | *EGFR* Del19 | DMSO | Osimertinib |
| 10 TH107 CDK6 | *EGFR* Del19 | DMSO | Osimertinib |
| *In vivo* models | Oncogenic Driver | Treatment | |
| 13 H1975 E.V. | *EGFR* L858R, T790M | Vehicle | Osimertinib |
| 14 H1975 CDK4 | *EGFR* L858R, T790M | Vehicle | Osimertinib |
| 15 H1975 CDK6 | *EGFR* L858R, T790M | Vehicle | Osimertinib |
| 16 PC9 E.V. | *EGFR* Del19 | Vehicle | Osimertinib |
| 17 PC9 CDK6 | *EGFR* Del19 | Vehicle | Osimertinib |
| 18 TH116 *CDK6* amp | *EGFR* L858R | Vehicle | Osimertinib |

Supplementary Table 3 – EGFR mutant NSCLC Foundation Medicine case's summary statistics for cell cycle gene (top) Copy Number (CN) alterations, concurrent CDK4 and CDK6 alterations (middle), and cell cycle gene CN changes concurrent with *CDK4* or *CDK6* amplifications (bottom)

| FM-LUAD | | | | |
|---|---|---|---|---|
| CC gene | CN Number of Cases Positive | Negative | % of EGFRmt cases 660 total | % of CC positive cases 357 total |
| *CDKN2A* | 150 | 510 | 22.7 | 42.0 |
| *CDKN2B* | 135 | 525 | 20.5 | 37.8 |
| *CDK4* | 52 | 608 | 7.9 | 14.6 |
| *CCNE1* | 33 | 627 | 5.0 | 9.2 |
| *RB1* | 28 | 632 | 4.2 | 7.8 |
| *CCND1* | 26 | 634 | 3.9 | 7.3 |
| *CDK6* | 19 | 641 | 2.9 | 5.3 |
| *CCND3* | 15 | 645 | 2.3 | 4.2 |
| *CDKN1A* | 7 | 653 | 1.1 | 2.0 |
| *CDKN2C* | 3 | 657 | 0.5 | 0.8 |
| *CCND2* | 3 | 657 | 0.5 | 0.8 |
| *CDKN1B* | 1 | 658 | 0.2 | 0.3 |

| | Concurrent CN | Del | SV |
|---|---|---|---|
| *CDK4* | 1 case | 0 | 6 cases |
| *CDK6* | | 0 | 6 cases |

| Concurrent CN | | | % | |
|---|---|---|---|---|
| | Number of Cases *CDK4* | *CDK6* | *CDK4* 52 total | *CDK6* 19 total |
| *CDKN2A* | 2 | 8 | 4 | 42 |
| *CDKN2B* | 2 | 7 | 4 | 37 |
| *CDK4* | 52 | 1 | 100 | 5 |
| *CCNE1* | 0 | 2 | 0 | 11 |
| *RB1* | 1 | 2 | 2 | 11 |
| *CCND1* | 2 | 1 | 4 | 5 |
| *CDK6* | 1 | 19 | 2 | 100 |
| *CCND3* | 0 | 1 | 0 | 5 |
| *CDKN1A* | 0 | 1 | 0 | 5 |
| *CDKN2C* | 0 | 0 | 0 | 0 |
| *CCND2* | 0 | 0 | 0 | 0 |
| *CDKN1B* | 0 | 0 | 0 | 0 |

Supplementary Table 3 (*continued*) – EGFR mutant NSCLC MSK-IMPACT GENIE case's summary statistics for cell cycle gene (top) Copy Number (CN) alterations, concurrent CDK4 and CDK6 alterations (middle), and cell cycle gene CN changes concurrent with *CDK4* or *CDK6* amplifications (bottom)

| MSK-GENIE-LUAD | | | | |
|---|---|---|---|---|
| CC gene | CN Number of Cases | | % of EGFRmt cases | % of CC positive cases |
| | Positive | Negative | 1983 total | 609 total |
| *CDKN2A* | 322 | 287 | 16.2 | 52.9 |
| *CDKN2B* | 301 | 308 | 15.2 | 49.4 |
| *CDK4* | 137 | 472 | 6.9 | 22.5 |
| *CCNE1* | 87 | 522 | 4.4 | 14.3 |
| *RB1* | 43 | 566 | 2.2 | 7.1 |
| *CCND1* | 42 | 567 | 2.1 | 6.9 |
| *CDK6* | 15 | 594 | 0.8 | 2.5 |
| *CCND3* | 25 | 584 | 1.3 | 4.1 |
| *CDKN1A* | 4 | 605 | 0.2 | 0.7 |
| *CDKN2C* | 6 | 603 | 0.3 | 1.0 |
| *CCND2* | 7 | 602 | 0.4 | 1.1 |
| *CDKN1B* | 16 | 593 | 0.8 | 2.6 |
| *E2F3* | 4 | 605 | 0.2 | 0.7 |

| | Concurrent CN | Del | SV |
|---|---|---|---|
| *CDK4* | 0 cases | 0 | 12 cases |
| *CDK6* | | 0 | 5 cases |

| Concurrent CN | | | % | |
|---|---|---|---|---|
| | Number of Cases | | | |
| | *CDK4* | *CDK6* | *CDK4* 137 total | *CDK6* 15 total |
| *CDKN2A* | 11 | 6 | 8 | 40 |
| *CDKN2B* | 11 | 5 | 8 | 33 |
| *CDK4* | 137 | 0 | 100 | 0 |
| *CCNE1* | 2 | 0 | 1 | 0 |
| *RB1* | 0 | 0 | 0 | 0 |
| *CCND1* | 8 | 1 | 6 | 7 |
| *CDK6* | 0 | 15 | 0 | 100 |
| *CCND3* | 8 | 0 | 6 | 0 |
| *CDKN1A* | 1 | 0 | 1 | 0 |
| *CDKN2C* | 0 | 0 | 0 | 0 |
| *CCND2* | 0 | 0 | 0 | 0 |
| *CDKN1B* | 1 | 0 | 1 | 0 |
| *E2F3* | 0 | 0 | 0 | 0 |
